# A Single-cell and Spatially Resolved Cell Atlas of Human Esophageal Squamous Cell Carcinoma

**DOI:** 10.1101/2025.06.01.657221

**Authors:** Yong Shi, Ke An, Yu Qi, Xinhan Zhang, Yueqin Wang, Xuran Zhang, Shaoxuan Zhou, Ouwen Li, Yanan Song, Jiayi Zhou, Yue Du, Mingyang Hou, Yun-Gui Yang, Quancheng Kan, Xin Tian

**Author notes:** Corresponding authors (Tian X), (Yang YG), (Kan Q). Equal contribution (Yong Shi, Ke An, Yu Qi, Xinhan Zhang).

## Abstract

Tumor heterogeneity and the suppressive microenvironment are key challenges that limit the effectiveness of cancer treatment. In this study, we systematically elucidated the molecular characteristics and mechanisms underlying the suppressive immune microenvironment via a combination of single-cell RNA sequencing, spatial transcriptomics, and metabolomics for a series of human esophageal squamous cell carcinoma (ESCC) and matched nontumor tissues. We found that *COL17A1*^+^ epithelial cells presented greater malignancy, characterized by triglyceride (TG) and phosphocholine (PC) accumulation. We also identified a tumor-specific *POSTN*^+^ fibroblast subgroup. We found a unique epithelial-fibroblast niche with low infiltration of effector immune cells and substantial enrichment of lipids, composed of *POSTN*^+^ fibroblasts and *COL17A1*^+^ epithelial cells, where their crosstalk contributed to tumor progression. We confirmed that the *INHBA*/*TP63* axis plays a key role in mediating the regulation of *COL17A1*^+^ tumor cells by *POSTN*^+^ fibroblasts. Our findings provided new insights into the characteristics of the tumor microenvironment and the crosstalk between tumor and fibroblasts, offering valuable multiomics data resources for elucidating tumor progression mechanisms.

## Introduction

Esophageal cancer is the seventh most prevalent malignancy and the sixth leading cause of tumor-associated death globally [1]. Esophageal squamous cell cancer (ESCC) accounts for ∼90% of these cases [2]. Patients with early-stage ESCC generally receive effective treatment after excision. However, after ESCC reaches an advanced stage or metastasizes, most patients succumb to the disease. Immune checkpoint blockade (ICB) strategies are used for treating advanced and metastatic diseases, including ESCC [3, 4]. Although ICB strategies have several benefits, their effectiveness is often limited by tumor microenvironment (TME) heterogeneity [5]. Therefore, the molecular characteristics of the TME need to be elucidated.

The combined analysis of single-cell data and spatial data not only provides transcriptional profiles of individual cells but also preserves their spatial information, offering significant advantages in investigating the heterogeneity of solid tumors and the features of the TME [6, 7]. Integrative analysis strategies have been widely used in various types of cancer, resolving the interactions of key cellular subpopulations associated with tumor progression, metastasis, and resistance to treatment [8–10]. Several studies on ESCC have focused on identifying critical protumor molecular events [11–13] and characterizing the distribution of immune and stromal cells in metastatic and nonmetastatic samples [14]. Therefore, further assessment of the cell interactions in the TME and how these interactions affect the spatial architecture of the microenvironment is essential.

Besides single-cell and spatial data, additional omics information is necessary to gain comprehensive insights into the TME. Disorders of cellular interaction and metabolism are two key determinants that significantly affect tumor immune evasion, invasion, metastasis, and resistance to therapy [15, 16]. The variations in metabolism are closely related to spatial heterogeneity [17]. The development of matrix-assisted laser desorption/ionization mass spectrometry imaging (MALDI-MSI) has provided a key method for examining the spatial features of metabolites in heterogeneous diseased tissues [18], thus enabling the delineation of the metabolic profile in the TME. Therefore, the integration of multiomics data may provide a comprehensive understanding of the mechanisms underlying tumor progression and reprogramming of the TME.

In this study, by integrating single-cell transcriptomics, spatial transcriptomics (ST), and spatial metabolomics (SM), we found important molecular events in the suppressive immune microenvironment. We also showed the heterogeneity and molecular features of tumor cells and fibroblasts. We identified a spatial structure composed of specific tumor cells and fibroblasts in ESCC characterized by a decrease in immune cell infiltration, emphasizing its importance in tumor immune evasion. We also systematically deciphered the mutual regulatory network and functions of tumor cells and fibroblasts within this structure. Overall, the findings of this study improved our understanding of the complexity of the TME in ESCC.

## Results

### Single-cell and spatial multiomics profiling of ESCC

To establish a single-cell and spatial multiomics atlas of ESCC, we included five ESCC patients (Table S1). The integrated analysis workflow is shown in Figure S1A. Single-cell RNA sequencing (scRNA-seq) was performed on five tumor tissues and three paired adjacent nontumor tissues. ST [19] and MALDI-MSI analyses [20] were conducted on consecutive sections from the five tumor tissue samples. A total of 92,526 cells were obtained from scRNA-seq. After quality control, 86,982 high-quality cells were retained, with an average sequencing depth of 1,243 genes per cell (Figure S1B and C). The data were integrated using the robust principal component analysis (RPCA) method, which effectively minimized batch effects (Figure S1D and E). 20 cell clusters were identified via unsupervised clustering (Figure S1F). Based on the mRNA expression of the known cell-specific signature genes, eight major cell clusters were annotated across tumor and adjacent nontumor tissues, including epithelial cells (*KRT18*, *EPCAM*, and *CD24*), endothelial cells (*PECAM1*, *CLDN5*, and *CD34*), fibroblasts (*DCN*, *MYL9*, and *COL3A1*), T/NK cells (*CD3E*, *CD3D*, and *CD8A*), B cells (*CD79A*, *JCHAIN*, and *IGHG1*), myeloid cells (*CD14*, *C1Q1*, and *C1QB*), mast cells (*TPSB2*, *TPSAB1*, and *CPA3*), and cycling immune cells (*MKI67*, *TOP2A*, and *PTPRC*) (**Figure 1**A and B, Figure S1G). The results of the cell type proportion analysis reflected the degree of variation in cell abundance among the ESCC samples, highlighting the heterogeneity and complexity of the interpatient tumor tissues (Figure 1C and D).

**Figure 1.**
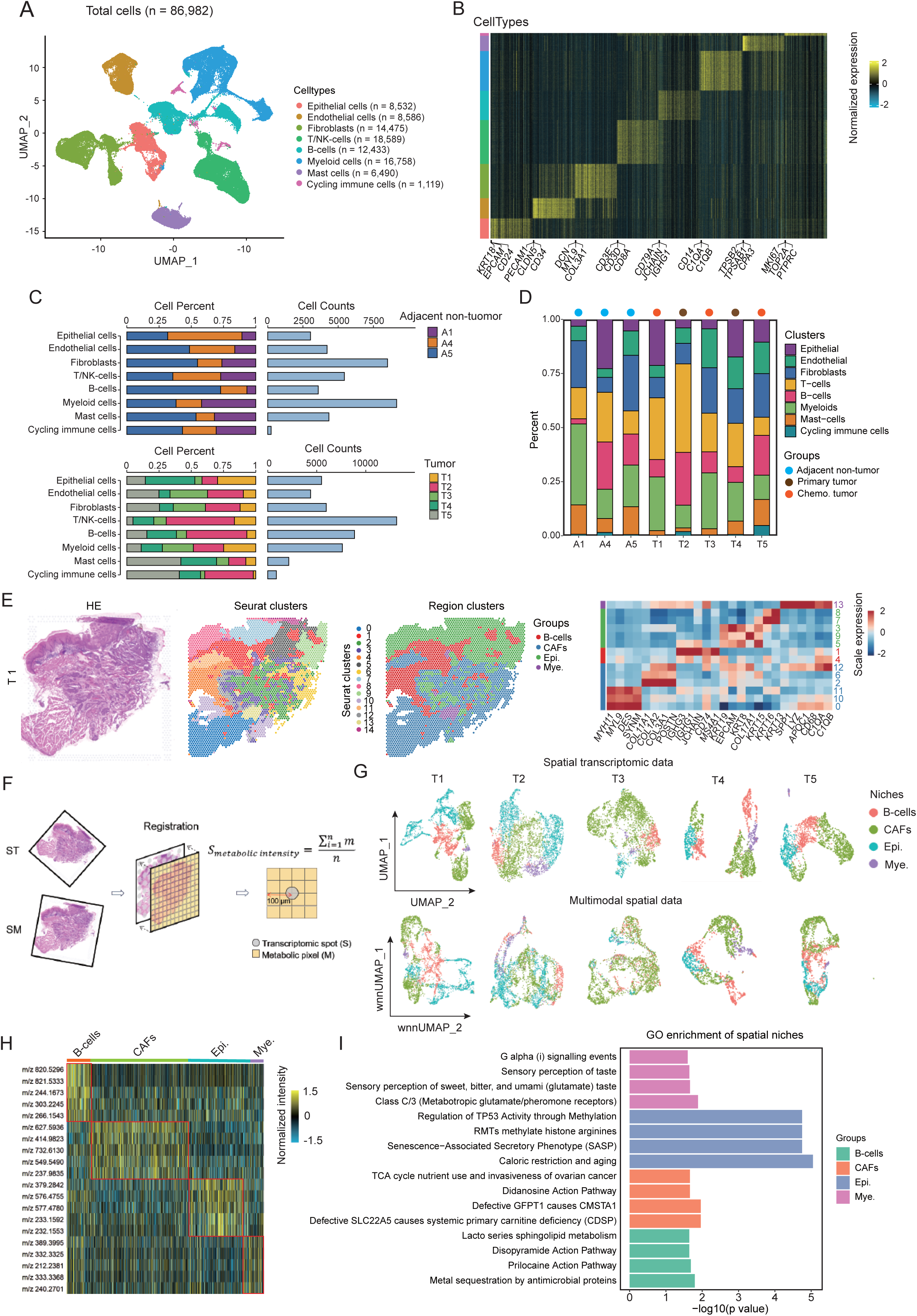
Single-cell and multiomics cell atlas in ESCC. **A.** UMAP plots of total cells labeled by main cell types from paired adjacent nontumor and tumor tissues in ESCC. **B.** Heatmap of the top 50 cell-type-specific genes for each major cell type in ESCC. **C.** Bar plots showing the proportion of eight cell types stratified by tumor sample, tissue type, and total cell count. **D.** Bar plots showing the cell proportion of major cell types across eight samples. **E.** Left: H&E staining of tissue sections; unbiased clustering of ST spots and four spatial niche clusters. Right: heatmap showing the expression of marker genes for four spatial niches including B cell, CAF, epithelial cell, myeloid. **F.** Integrated analysis workflow of ST and MALDI-MSI data. **G.** UMAP plots of ST data and WNN-UMAP plots of multimodal data both labeled by four spatial niches. **H.** The heatmap displaying the top 10 differentially abundant metabolites for each spatial niche. **I.** Bar plots showing the top four GO terms enriched for niche-specific metabolites. UMAP, uniform manifold approximation and projection; H&E, hematoxylin and eosin; CAF, cancer associated fibroblast; ST, spatial transcriptomic; MALDI-MSI, matrix-assisted laser desorption/ionization mass spectrometry imaging; WNN, weighted nearest neighbor; GO, gene ontology.

For ST data, we obtained valid transcriptomic data from 3653, 3790, 2856, 1710, and 2169 spots in five samples, with median depths of 8817, 8024, 1968, 9414, and 3219 unique molecular identifiers (UMI) counts per spot and 3130, 2877, 1071, 3320, and 1473 genes per spot, respectively (Figure S2A and B). Integrated analysis of ST and scRNA-seq revealed spatially restricted distribution patterns for distinct cell types (Figure S2C). By conducting unbiased clustering via Seurat, the spots were categorized into 15, 9, 9, 7, and 9 clusters for the five sections, respectively (Figure 1E, Figure S2D). Based on marker gene expression from these spatial clusters, we annotated all sections into four regions: epithelial cell, B cell, cancer associated fibroblast (CAF), and myeloid niches (Figure 1E, Figure S2D).

For MALDI-MSI analyses, 1474 metabolites were detected across five tumor samples. To further integrate ST with SM data, we adopted a novel workflow involving two major steps: (1) alignment of ST and SM images and (2) calculation of the metabolic intensity for each spatial spot (Figure 1F). We combined ST and SM data to create multimodal data and mapped it to the weighted nearest neighbor-UMAP (WNN-UMAP) space (Figure 1F). The four spatial niches also generated separate clusters in the WNN-UMAP space, confirming the effectiveness and robustness of our integration methods (Figure 1G). By conducting differentially abundant metabolite analysis, we identified spatial niche-specific metabolites (Figure 1H). Further functional enrichment analysis revealed that metabolomics varied across different spatial niches, such as significant enrichment of sphingolipid metabolism in the B cell niche, RMTs that methylate histone arginine in the epithelial cell niche, the TCA cycle in the fibroblast niche, and metabotropic glutamate in the myeloid niche (Figure 1I). To summarize, we constructed a single-cell and spatial multiomics cell atlas of ESCC.

### The immune landscape of ESCC

The occurrence and development of tumors are closely related to the TME [21]. To gain insights into the spatial organization of the immune microenvironment in ESCC, we first conducted subpopulation clustering for immune cells. After unbiased clustering, total immune cells could be split into five major compartments and further annotated into 19 distinct subpopulations, each distinguished by specific marker genes (Figure S3A–C). The expression of known key immune genes was evaluated across immune cell subtypes (**Figure 2**A). Three CD8^+^ T-cell subsets (*CXCL13*^+^ CD8^+^ T cells, *GNLY*^+^ CD8^+^ T cells, and cycling CD8^+^ T cells) expressed high levels of immune effector molecules and exhibited high expression of immune inhibitory molecules, reflecting immune exhaustion in the TME (Figure 2A). Both *FGFBP2*^+^ NK and *AREG*^+^ NK cells presented high expression of immune effector molecules. However, *FGFBP2*^+^ NK cells expressed inhibitory molecules at lower levels and had good prognostic value, suggesting that they may be in an immune-activated state (Figure 2A and B).

**Figure 2.**
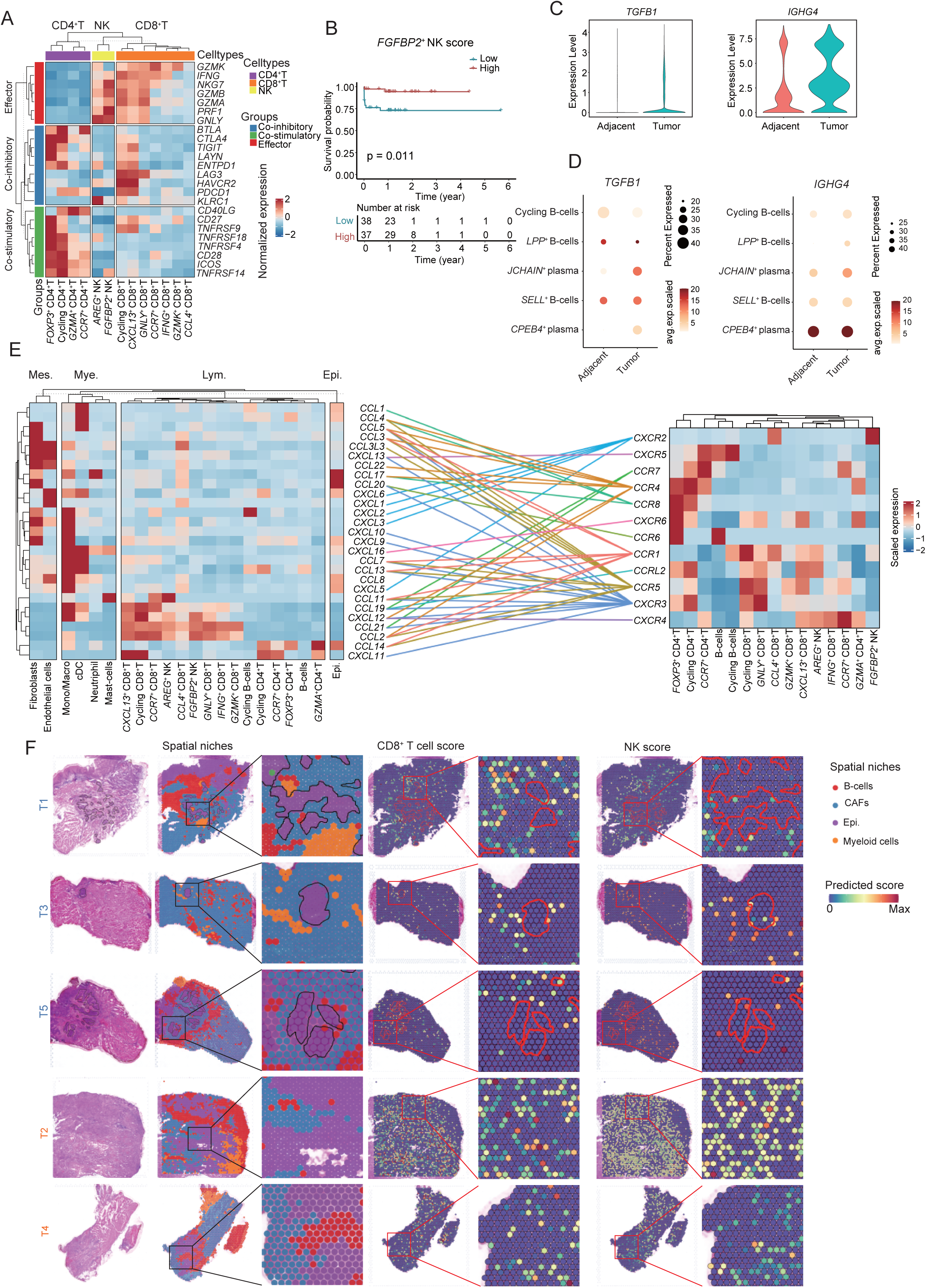
The immune landscape of ESCC. **A**. Scaled mean normalized mRNA expression heatmap of genes related to cytotoxicity, co-inhibitory, and co-stimulatory molecules across T cell and NK subsets in scRNA-seq data. **B.** Kaplan–Meier survival curve for *FGFBP2*^+^ NK cells from a TCGA cohort of 75 ESCC patients. **C.** Violin plots of *TGFB1* and *IGHG4* comparing B cells in adjacent versus tumor tissues based on scRNA-seq data. **D.** Dot plots showing the mRNA expression of *TGFB1* and *IGHG4* between nontumor and tumor tissues among B cell subpopulations. **E.** Left: scaled mean normalized mRNA expression heatmap of chemokine genes in distinct cell types from single-cell sequencing data. Right: scaled mean normalized mRNA expression heatmap of chemokine receptor genes in distinct lymphocyte cell types from scRNA-seq data. Chemokines and matching receptors were linked by color lines. **F.** Left: H&E staining and spatial clusters with annotated tumor region (solid lines) across five tumor samples. Right: spatial feature plots showing the enrichment score of CD8^+^ T and NK cells across five tumor samples. A p-value < 0.05 is considered as a statistically significant difference. Statistical significance was assessed using a two-sided log-rank test in (B). scRNA-seq, single-cell RNA sequencing; TCGA, the cancer genome atlas.

Our multiomics data revealed that B cells constitute a major component of the TME (Figure S2D, Figure S3A), suggesting that B cells may play key roles in reshaping the TME. Another study reported that an important B-cell subpopulation, known as regulatory B cells (Bregs), can weaken the immune response to tumors by secreting *TGFB1 or IGHG4* [22, 23]. Our data revealed a significant increase in the expression of *TGFB1* and *IGHG4* in B cells within tumor tissues (Figure 2C). Compared to that in adjacent normal tissues, the expression of *TGFB1* was significantly greater in *CPEB4*^+^ plasma cells, and *JCHAIN*^+^ plasma cells within tumor tissues, and *IGHG4* expression was markedly elevated in *LPP*^+^ B cells and *JCHAIN*^+^ plasma cells in tumor tissues (Figure 2D). Spatial analysis also revealed high mRNA expression of *IGHG4* and *TGFB1* in the B-cell niche (Figure S3D). Additionally, several classic immunosuppressive molecules also occur in B cells (Figure S3E). Hence, our findings suggest that B cells are involved in the generation of the inhibitory microenvironment.

Potential mechanisms underlying the recruitment of tumor-infiltrating lymphocytes were next assessed by analyzing chemokine and receptor expression in the scRNA-seq data (Figure 2E). Half of the detected chemokine receptors were expressed predominantly in regulatory T cells (Tregs) (Figure 2E). Most chemokine recruited Tregs were highly expressed in fibroblasts, endothelial cells, mono/macrophages, and cDCs, suggesting that these cells are responsible for recruiting Tregs (Figure 2E). *FGFBP2*^+^ NK cells, which are important antitumor effector cells, expressed only one chemokine receptor from our detected chemokine pool, implying insufficient recruitment of *FGFBP2*^+^ NK cells in the TME and limiting their antitumor effects (Figure 2E).

Major antitumor immune cell types, including CD8^+^ T cells and NK cells, were mapped to the ST data. The results of the quantitative analysis revealed a decrease in the colocalization of tumor cells with CD8^+^ T or NK cells in tumor sample 1, 3, and 5 (Figure S3F and G). A unique spatial barrier was formed in tumor sample 1/3/5, where CAFs enveloped tumor cells (Figure 2F). Therefore, we speculated that the reduction in immune cell infiltration may be related to the formation of spatial barriers. In samples with barrier formation, CD8^+^ T and NK cells were restricted to the fibroblast-rich regions near the tumor, whereas in samples without barrier formation, they were distributed within the tumor region (Figure 2F). We also found that immune effector molecules presented a lower abundance in the tumor region, generating a CAF barrier (Figure S3H). By integrating ST and SM data, we found that the barrier region was significantly enriched with lipid metabolites (Figure S4A and B). Consistent with these findings, ST data revealed significant enrichment of the lipid synthesis-related genes *ACACA* [24], *ACLY* [25], and *FASN* [26] in the barrier region (Figure S4C). Some studies have shown that lipids have cytotoxic effects on immune cells [27–29]. These findings suggest that the enrichment of lipid metabolites in barrier regions is a significant factor contributing to the decrease in the infiltration of immune cells. These results highlight multiple cell types involved in immunosuppressive mechanisms, including (1) the recruitment of Tregs, (2) B cells that exhibit Bregs characteristics, (3) exhausted T cells, (4) insufficient recruitment of *FGFBP2*^+^ NK cells, and (5) the formation of a spatial barrier niche with a decrease in the number of effector immune cells and an increase in the enrichment of lipids.

### Two recurring tumor epithelial subpopulations

After unsupervised clustering, all epithelial cells were divided into nine subgroups (Figure S5A). Further copy number variation (CNV) analysis revealed that subgroups 1, 7, and 8 had the lowest CNV scores, suggesting that these cells were nonmalignant (Figure S5B and C). To eliminate the effect of nonmalignant cells, we excluded these cells from the heterogeneity analysis. After reclustering total malignant epithelial cells, two epithelial clusters were generated, each expressing representative genes, including *LCN2^+^* cells (marked by the genes *LCN2*, *SLPI*, and *S100P*) and *COL17A1*^+^ cells (marked by the genes *COL17A1*, *EGFR*, and *KRT15*) (Figure 3A, Figure S5D). Epithelial cells from the external dataset could also be classified into two subpopulations based on the top 13 cell-type-specific genes of each epithelial subpopulation (Figure 3C, Figure S5E). Functional analysis revealed that *LCN2^+^*epithelial cells were enriched in pathways related to the maturation of squamous epithelial cells, such as epidermis differentiation, development, and keratinization, whereas *COL17A1^+^* epithelial cells were enriched in extracellular matrix (ECM) formation (Figure S5F). *COL17A1*^+^ epithelial cells preferentially occurred in tumor tissues, indicating increased malignancy (Figure S5G). *COL17A1*^+^ epithelial cells presented higher epithelial-mesenchymal transition (EMT) scores (Figure 3D) and a greater proportion of proliferating cells (Figure 3E and F); these cells were associated with poorer patient prognosis (Figure S5H). These cells also presented lower expression levels of squamous epithelial differentiation-related genes, such as *KRT16*, *KRT13*, *KRT6B*, *KRT6C*, *MUC21*, *MUC4*, *CXCL17*, *S100A8*, and *S100A9* [30, 31], further indicating that *COL17A1*^+^ epithelial cells were more malignant and in a dedifferentiated state (Figure 3G). Subsequent cell trajectory analysis confirmed the dedifferentiated state of *COL17A1*^+^ epithelial cells (Figure 3A and B). These results indicate that *COL17A1*^+^ epithelial cells may be more aggressive and malignant and that *LCN2*^+^ cells are tumor progenitor cells.

**Figure 3.**
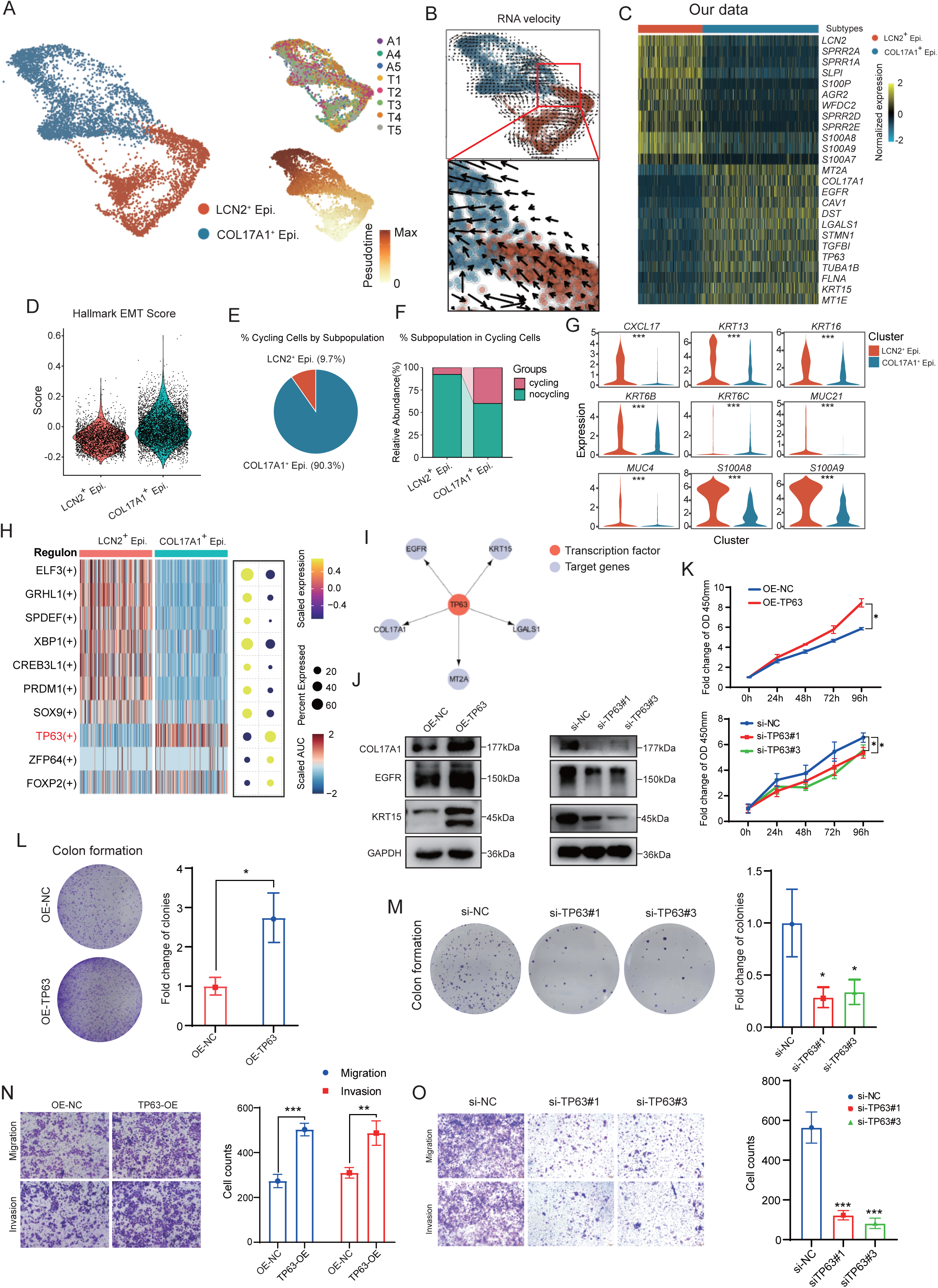
Identification of two recurrent subpopulations in epithelial cells. **A.** UMAP plots showing epithelial cells labeled by two subpopulations, samples, and pseudo-time, respectively. **B.** Single-cell trajectory inference of epithelial cells using scVelo. **C.** Heatmap showing the top 13 highly expressed genes for each epithelial subset. **D.** Violin plots showing the EMT pathway activity scores in epithelial subpopulations. **E.** Pie plots showing the percentage of epithelial subpopulations within cycling cells. **F.** Bar plots showing the percentage of cycling cells within epithelial subpopulations. **G.** Violin plots showing the expression of epidermis maturation-related genes including *KRT16*, *KRT13*, *KRT6B*, *KRT6C*, *MUC21*, *MUC4*, *CXCL17*, *S100A9*, *S100A8* across epithelial cell subpopulations. **H.** Heatmap and dot plots showing the specific TF regulon AUC scores and mRNA expression in epithelial subpopulations. **I.** Regulatory network of TF *TP63* and its downstream target genes. **J.** WB analysis showing the changes in protein expression levels of *KRT15*, *COL17A1*, and *EGFR* in KYSE-150 cells following *TP63* overexpression or knockdown. **K.** Cell viability was measured at 0, 24, 48, 72, and 96 h after *TP63* overexpression or knockdown in KYSE-150 cells using a CCK-8 assay (n = 3). Quantification of colony formation efficiency in KYSE-150 cells following *TP63* overexpression (**L**) or knockdown (**M**) (n = 3). Representative migration and invasion images of KYSE-150 cells in Transwell chambers at 24 and 48 h after *TP63* overexpression (**N**) or knockdown (**O**) (n = 3). Scale bar = 100 μm. *, *P* < 0.05; **, *P* < 0.01; ***, *P* < 0.001. Statistical significance was calculated by Student’s *t*-test, mean ± SD. EMT, epithelial-mesenchymal transition; TF, transcription factor; AUC, area under curve; CCK-8, Cell Counting Kit-8; WB, western blot; SD, standard deviation.

Transcription factor (TF) is key determinants of cell fate. Therefore, we performed single-cell regulon network inference using the SCENIC software [32]. (Figure 3H). We found that the TF *TP63* was most highly expressed in *COL17A1*^+^ epithelial cells, and its target genes, including *COL17A1*, *EGFR*, *MT2A*, *KRT15*, and *LGALS1*, were characteristic genes of *COL17A1*^+^ epithelial cells (Figure 3C and I). These findings suggest that *TP63* may be closely related to the lineage differentiation of *COL17A1*^+^ epithelial cells. To test this hypothesis, we conducted in *vitro* experiments in ESCC cell lines. The results revealed that the overexpression of *TP63* promoted the expression of the mRNAs of these characteristic genes, whereas knocking down *TP63* reduced their mRNA levels (Figure S6A–D). We also found that the overexpression of *TP63* increased the protein levels of *COL17A1*, *EGFR*, and *KRT15* (Figure 3J). These findings suggest that *TP63* is a crucial determinant of the fate of *COL17A1*^+^ epithelial cells.

To validate the function of *TP63*, we examined changes in cell function after overexpressing or knocking down *TP63* in ESCC cells. The results indicated that overexpressing *TP63* significantly increased tumor cell viability, clonogenic ability, and migration and invasion capabilities, whereas knocking down *TP63* had the opposite effect (Figure 3K–O). In sum, we identified two epithelial subpopulations in ESCC tissues, with *COL17A1*^+^ epithelial cells exhibiting increased malignancy, and the TF *TP63* plays a key role in maintaining the malignant phenotype of *COL17A1*^+^ epithelial cells.

### Spatial heterogeneity of epithelial cells

To delineate the spatial heterogeneity of epithelial cells, two epithelial subpopulations were mapped to ST data. This analysis revealed that distinct epithelial subpopulation had a regionally restricted distribution tendency (**Figure 4**A). The marker genes *LCN2* and *COL17A1* also showed spatial patterns similar to those of the deconvolution scores (Figure 4A). Immunofluorescence experiments confirmed these distinct spatial expression features (Figure 4B and C). We also found that the spatial microenvironment correlated with transcriptomic alterations in *COL17A1*^+^ epithelial cells. For example, the distribution patterns of *COL17A1*^+^ epithelial cells include two distinct niches: those adjacent to B cells and those intertwined with fibroblasts, as described in tumor sample 1 (Figure 4D). In *COL17A1*^+^ epithelial cells from the fibroblast niche, the upregulated genes were enriched in the EMT-related pathway, whereas those from the B-cell niche were associated with the reactive oxygen species (ROS) pathway, reflecting the plasticity of tumor cell states driven by spatial organization (Figure 4E, Figure S7A). The results of the CNV analysis revealed that spatial factors could drive tumors to form distinct subclonal populations (Figure S7B and C), suggesting that spatial variables are correlated with transcriptomic plasticity and affect genomic variation in tumor cells.

**Figure 4.**
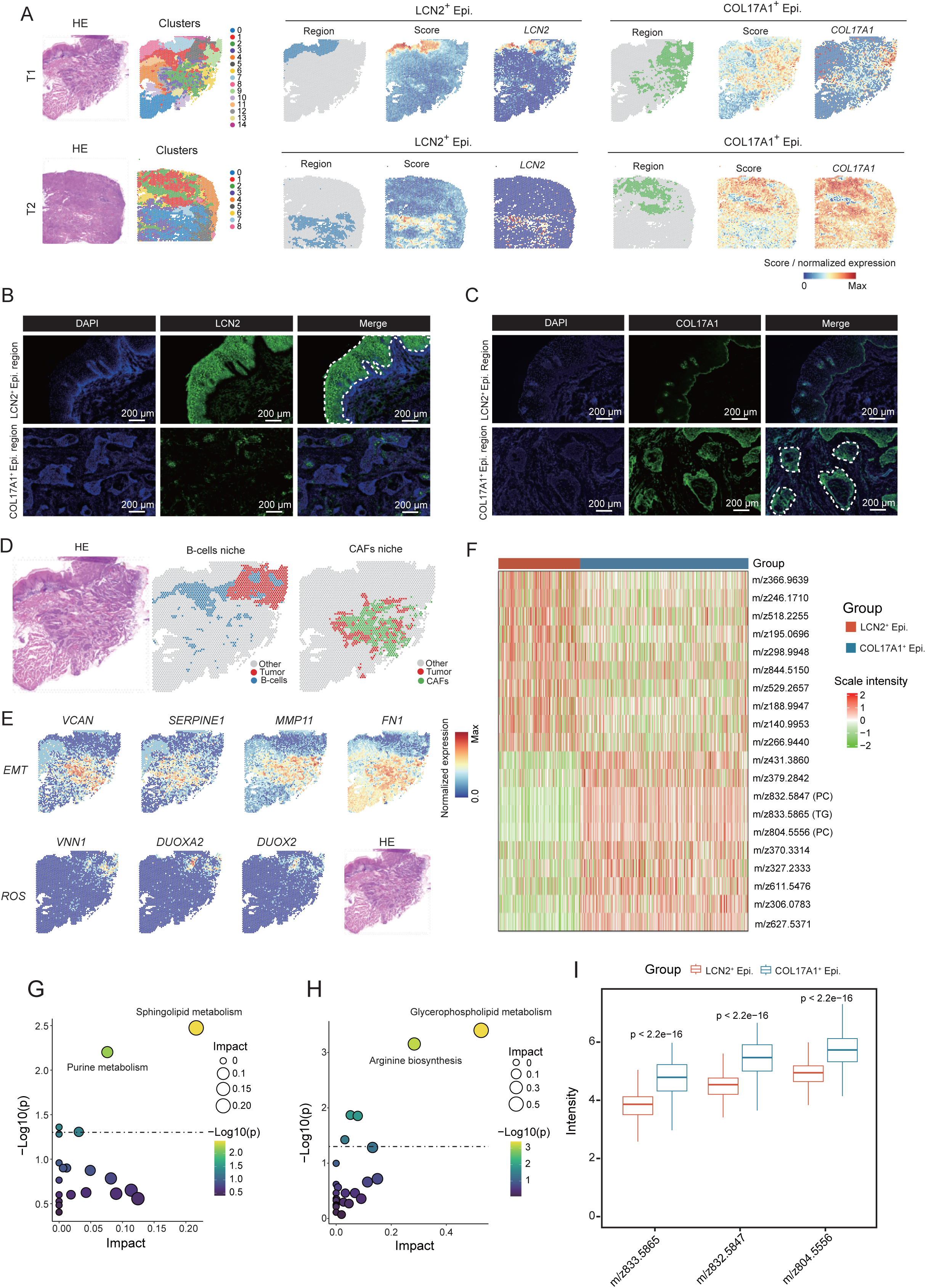
Spatial characteristics of epithelial subpopulations. **A.** Left: H&E-stained tissue sections, followed by ST spots unbiased clustering. Right: spatial feature plots showing epithelial subpopulation region, epithelial scores, and the expression of marker genes including *LCN2* and *COL17A1* in tissue sections for tumor sample 1 and 2. IF staining of *LCN2* (**B**) and *COL17A1* (**C**) in *LCN2*^+^ and *COL17A1*^+^ epithelial regions from tumor sample 1. Scale bar = 200μm. **D.** Left: H&E staining of tissue section for tumor sample 1. Right: spatial spots of *COL17A1*^+^ epithelial cells in B cell or CAF niches. **E.** Spatial feature plots of selected genes including *VCAN*, *SERPINE1*, *MMP11*, *FN1*, *VNN1*, *DUOXA2*, *DUOX2* in tumor sample 1. **F.** Heatmap of the top 10 differentially enriched metabolites for each epithelial cell subpopulation. **G.** Dot plots showing the enriched pathways of upregulated metabolites in *LCN2*^+^ epithelial cells. **H.** Dot plots showing the enriched pathways of upregulated metabolites in *COL17A1*^+^ epithelial cells. **I.** Box plots showing the enrichment levels of PC (m/z 832.5847, m/z 804.5556) and TG (m/z 833.5865) in epithelial cell subpopulations. ***, *P* < 0.001. Statistical significance was calculated by wilcoxon test, mean ± SD. IF, Immunofluorescence; PC, phosphocholine; TG, triglyceride.

To evaluate the metabolic heterogeneity of epithelial cells, we used a deconvolution method to determine the abundance and spatial distribution of the two epithelial cell types (Figure S7D–E). We then integrated SM data to identify their metabolic profiles. Differential analysis revealed 155 significantly increased metabolites in the *COL17A1*^+^ tumor region and 400 significantly increased metabolites in the *LCN2^+^* tumor region (Figure S7F). The top 10 differentially enriched metabolites are shown in Figure 4F. Further functional enrichment analysis revealed that *LCN2*^+^ epithelial cells were enriched in sphingolipid metabolism and purine metabolism, whereas *COL17A1*^+^ epithelial cells were enriched in arginine biosynthesis and glycerophospholipid metabolism (Figure 4G and H). The procarcinogenic metabolites triglyceride (TG) and phosphocholine (PC) significantly accumulated in the *COL17A1*^+^ tumor region, reflecting greater lipid accumulation in *COL17A1*^+^ epithelial cells [33, 34] (Figure 4I). W found that the mRNA levels of diacylglycerol-acyltransferase 1 (*DGAT1*) and lysophosphatidylcholine acyltransferase 1 (*LPCAT1*), which are related to the synthesis of TG and PC, were also high in *COL17A1*^+^ epithelial cells, as indicated by the scRNA-seq and ST data (Figure S7G–I). Overall, we found that the two types of epithelial cells occupied distinct spatial niches and that spatial factors were closely associated with the transcriptomic and genomic plasticity of tumors. We also determined their different metabolomic characteristics.

### *POSTN*^+^ fibroblasts are associated with tumor progression

To investigate the heterogeneity and functionality of fibroblasts in ESCC, we first eliminated batch effects across samples and conducted clustering analysis (Figure S8A and B). The results indicated that based on the expression levels of marker genes, fibroblasts could be categorized into seven subpopulations: *ACTG2*^+^ fibroblasts, *CD36*^+^ fibroblasts, *SORBS2*^+^ fibroblasts, *CFD*^+^ fibroblasts, *GFRA3*^+^ fibroblasts, *POSTN*^+^ fibroblasts, and unidentified fibroblasts (**Figure 5**A and B). Cell proportion analysis revealed that the abundance of fibroblast subgroups differed among tumor samples, reflecting the heterogeneity of the TME (Figure S8C). Functional enrichment analysis indicated that *ACTG2*^+^ fibroblasts were associated with oxidative phosphorylation and energy derivation by oxidation, *CD36*^+^ and *SORBS2*^+^ fibroblasts were associated with vascular processes in the circulatory system and muscle system processes, *CFD*^+^ fibroblasts were involved in immune regulatory functions, *GFRA3*^+^ fibroblasts were connected to axon development, and *POSTN*^+^ fibroblasts were related to the organization of extracellular structures (Figure S8D). We found that *GFRA3*^+^ and *POSTN*^+^ fibroblasts, especially *POSTN*^+^ fibroblasts, were preferentially distributed within tumor tissues, whereas other types of fibroblasts were predominantly located in adjacent nontumor tissues, indicating that they may play a key role in tumor progression (Figure 5C). Further large-scale analysis demonstrated that the enrichment of *POSTN*^+^ fibroblasts was greater in tumor tissues than in normal tissues and was associated with poor patient prognosis (Figure 5D and E).

**Figure 5.**
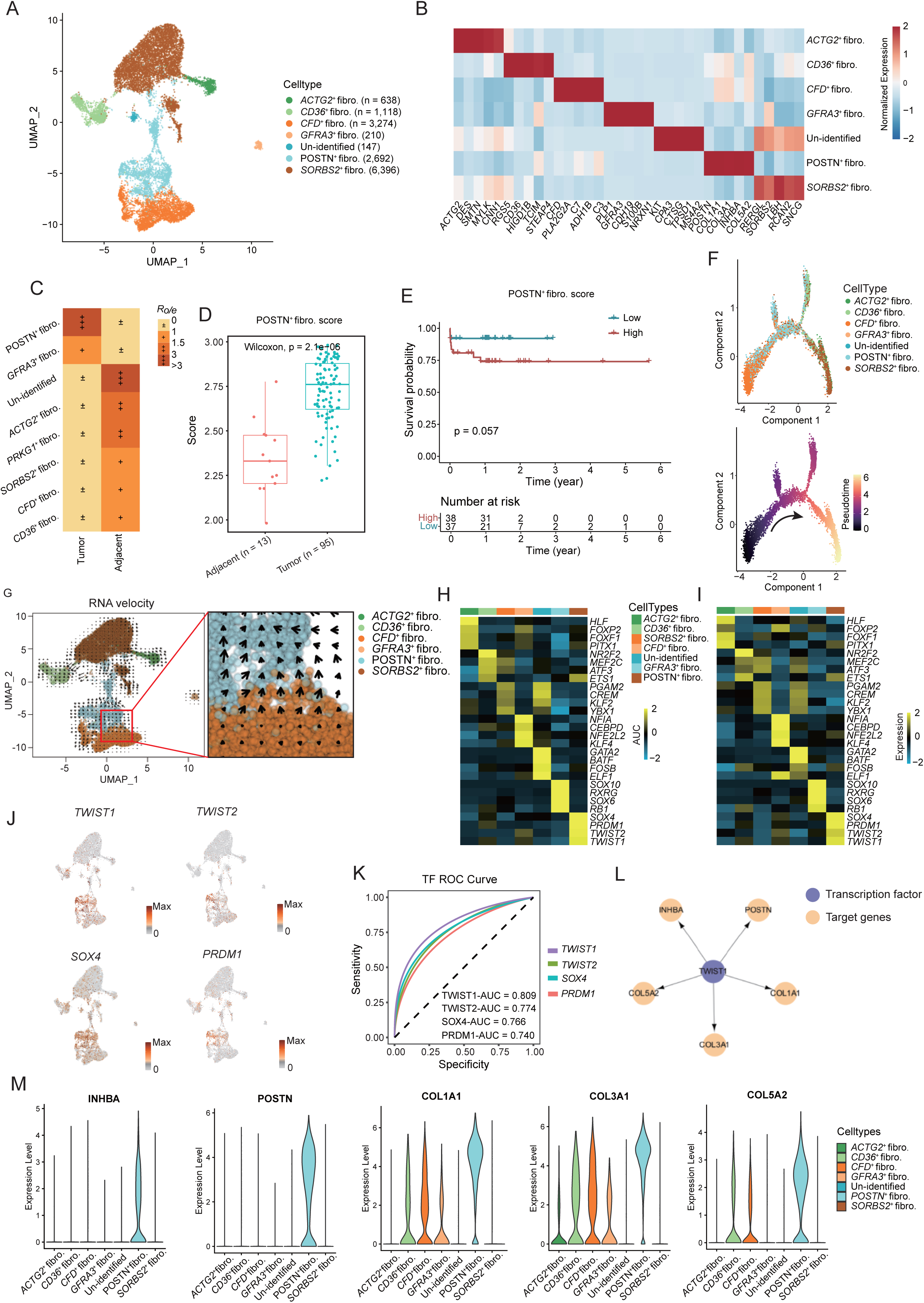
Analysis of fibroblast heterogeneity. **A.** UMAP plots showing total fibroblasts labeled by seven fibroblast subpopulations. **B.** Heatmap showing the mRNA expression of the top five marker genes for seven fibroblast subpopulations. **C.** Comparison of the tissue preference distribution among seven fibroblast subpopulations in paired samples (n = 3). The Ro/e value indicates tissue preference, with Ro/e > 1 suggesting enrichment in tumor tissues and Ro/e < 1 indicating preference for adjacent notumor tissues. **D.** Box plots showing the enrichment scores of *POSTN*^+^ fibroblasts between tumor (n = 95) and normal tissues (n = 13) in ESCC dataset from TCGA database. **E.** Survival curve of *POSTN*^+^ fibroblast score in 75 ESCC samples. **F.** UMAP plots of cell trajectory analysis labeled by fibroblast subpopulations and pseudo-time. **G.** Trajectory analysis of fibroblasts using scVelo. Heatmap showing the AUC scores (**H**) and mRNA expression (**I**) of the top four TF regulons for each fibroblast subpopulation. **J.** Feature plots of TF including *TWIST1*, *SOX4*, *TWIST2*, *PRDM1* in fibroblasts. **K.** ROC curve analysis showing the specificity of TF *TWIST1*, *TWIST2*, *SOX4*, and *PRDM1* in *POSTN*^+^ fibroblasts. **L.** Regulatory network of TF *TWIST1* and its selected target genes. **M.** Violin plots showing the mRNA expression of *INHBA*, *POSTN*, *COL1A1*, *COL3A1*, *COL5A2* among fibroblast subpopulations. *P* < 0.05 (two-sided log-rank test) is considered as a statistically significant difference. ROC, receiver operating characteristic curve.

We next performed cell trajectory analysis on these fibroblast subpopulations to determine the hierarchies of fibroblast differentiation. *POSTN*^+^ fibroblasts were positioned behind *CFD*^+^ fibroblasts in the inferred cell trajectory, suggesting that they probably originated from *CFD*^+^ fibroblasts (Figure 5F and G). Additionally, single-cell regulatory network inference was performed to identify subpopulation-specific TFs, which included the genes *TWIST1*, *TWIST2*, *SOX4*, and *PRDM1*, as TFs potentially controlling the fate of *POSTN*^+^ fibroblasts (Figure 5H). The mRNA expression of these TFs was also greater in *POSTN*^+^ fibroblasts than in other cells (Figure 5I and J). ROC curve analysis revealed that *TWIST1* exhibited the highest specificity for *POSTN*^+^ fibroblasts (Figure 5K). *TWIST1* downstream genes, including *POSTN*, *INHBA*, *COL1A1*, *COL3A1*, and *COL5A2*, were identified as characteristic genes of *POSTN*^+^ fibroblasts (Figure 5L and M). Thus, we selected *TWIST1* for experimental validation. Overexpressing *TWIST1* in the BJ cell significantly increased the mRNA expression of these genes, whereas knocking down *TWIST1* produced the opposite effect (Figure S8E–H). These findings indicate that *TWIST1* is a key determinant of the fate of *POSTN*^+^ fibroblasts. To summarize, we identified an important protumor fibroblast subgroup, known as *POSTN*^+^ fibroblasts, and found that *TWIST1* is a key TF that regulates the fate of these cells.

### *POSTN*^+^ fibroblasts and *COL17A1*^+^ epithelial cells are the dominant cell types in the immune barrier region

We found that a CAF barrier is formed in ESCC (Figure 2F). To decipher how the structure was organized, we first extracted the spots in the CAF barrier and calculated the enrichment levels of tumor cells and fibroblasts via spatial deconvolution in Seurat. We found that *POSTN*^+^ fibroblasts and *COL17A1*^+^ epithelial cells were the predominant cell types in the CAF barrier (**Figure 6**A and B). Cell neighborhood detection indicated that *POSTN*^+^ fibroblasts and *COL17A1*^+^ epithelial cells had the greatest number of colocalized spots in the barrier region, reflecting their spatial proximity (Figure 6C and D). To confirm our findings, the distribution patterns of *POSTN* and *COL17A1* in the barrier region were determined through immunofluorescence experiments. *POSTN* surrounded the tumor nests, forming a barrier structure, while *COL17A1* was highly expressed in the tumor nests (Figure 6E). We also found that the marker genes for CD8^+^ T cells or NK cells, *CD8A* and *CD16*, were predominantly distributed outside the *POSTN* barrier (Figure 6F). Our findings revealed that the immune barrier region was significantly enriched in lipid metabolites (Figure S4B). Therefore, we further investigated which cell type contributes to lipid enrichment in the barrier region by comparing the enrichment levels of lipid synthesis-related genes. Our results revealed that *COL17A1*^+^ epithelial cells had higher lipid synthesis-related gene signature scores, suggesting that they were major contributors to lipid enrichment in the barrier region (Figure 6G). These results revealed that the immune barrier primarily consisted of *POSTN*^+^ fibroblasts, which may shield immune surveillance in *COL17A1*^+^ epithelial cells. Additionally, *COL17A1*^+^ epithelial cells may further impair the activity of immune cells by enhancing lipid metabolism.

**Figure 6.**
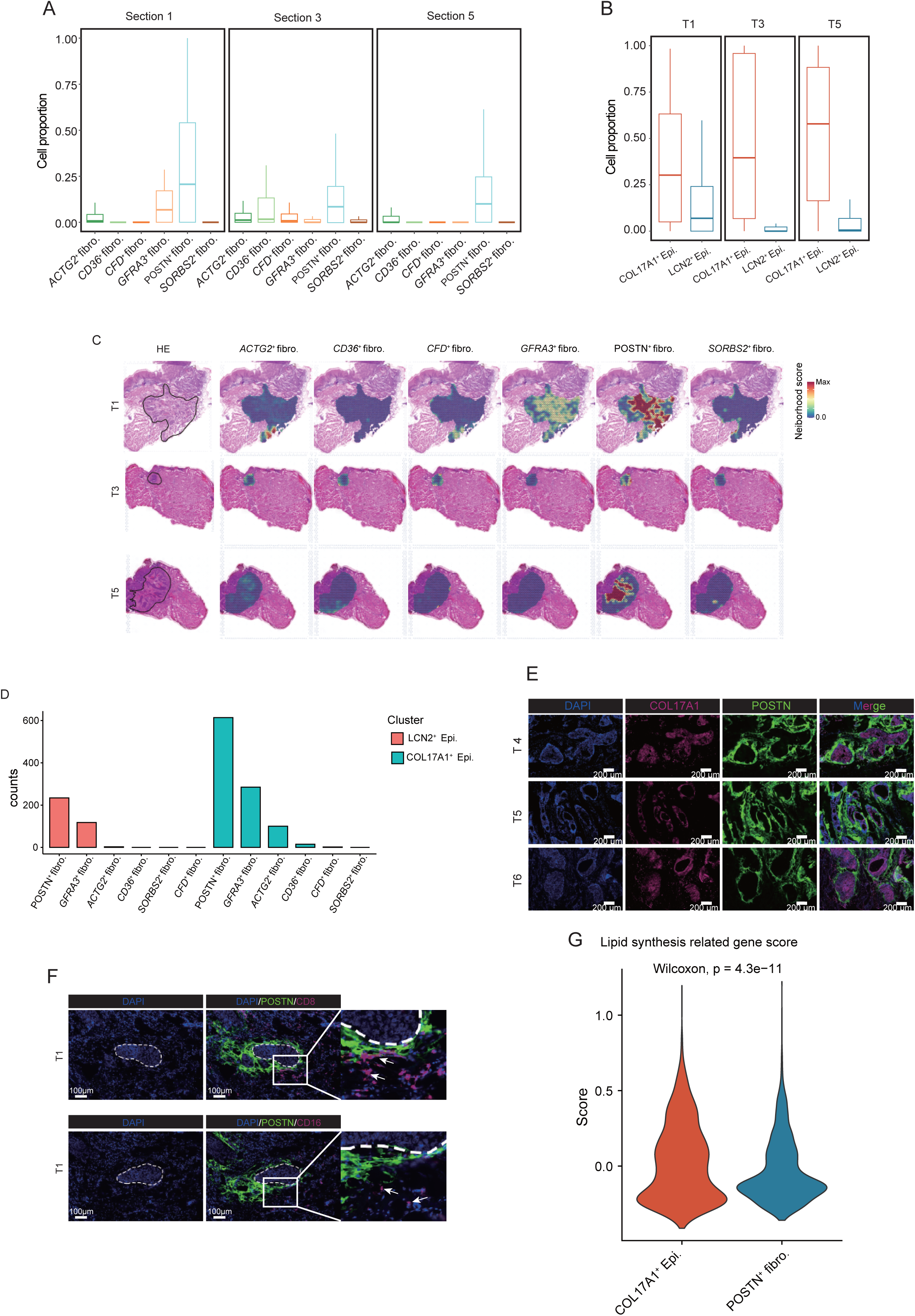
Identification of the components of the barrier niche. **A.** Box plots showing the enrichment scores of seven fibroblast subpopulations in annotated CAF barrier regions across tumor sample 1, 3, and 5. **B.** Box plots showing the enrichment scores of two epithelial subpopulations in annotated CAF barrier regions across tumor sample 1, 3, and 5. **C.** Left: H&E staining with annotated CAF barrier regions (solid lines) in tumor sample 1, 3, and 5. Right: spatial feature plots showing the neighborhood scores of fibroblast subpopulations within spatial barrier niches. **D.** bar plots showing the number of neighboring spots between epithelial and fibroblast subpopulations. **E.** Multi-color IF of *COL17A1* and *POSTN*. Scale bar = 200 μm. **F.** Representative multi-color IF staining of *POSTN*, *CD8*, and *CD16* in the immune barrier regions from tumor sample 1. *POSTN* represented fibroblasts, *CD8* represented CD8^+^ T cells, *CD16* represented NK cells. Scale bar = 100 μm. **G.** Violin plots comparing the lipid synthesis-related gene signature (*ACACA*, *ACLY*, *FASN*) scores between *COL17A1*^+^ epithelial cells and *POSTN*^+^ fibroblasts.

### *POSTN*^+^ fibroblast and *COL17A1*^+^ epithelial cell interactions contribute to tumor progression and immune barrier formation

To investigate the underlying mechanisms that contribute to the formation of this unique epithelial-fibroblast niche, we integrated scRNA-seq and ST data to resolve signaling communication between the two cell types. Based on the integrated ligand-receptor (L-R) pair pool from public datasets, we identified interactions between *POSTN*^+^ fibroblasts and *COL17A1*^+^ epithelial cells using the CellPhoneDB software. We detected a greater number of ligand-receptor pairs between *POSTN*^+^ fibroblasts and *COL17A1*^+^ epithelial cells than between other cell type pairs, suggesting a robust interaction between the two cell types (**Figure 7**A and B). Consistent with the spatial niches of *POSTN*^+^ fibroblasts and *COL17A1*^+^ epithelial cells, prominent *COL17A1*^+^ epithelial cell signaling to *POSTN*^+^ fibroblasts was mediated by several L-R pairs, such as *TGFB1*-*TGFBR1* and *BMP2*-*BMPR2* (Figure 7C, Figure S9A). In contrast, *POSTN*^+^ fibroblasts may modulate *COL17A1*^+^ epithelial cells through L-R pairs such as *COL3A1*-*CD44*, *BMP8A*-*CRABP2*, and *PLAU*-*ADORA2B* (Figure 7D, Figure S9B). *CFD*^+^ fibroblasts (a precursor of *POSTN*^+^ fibroblasts) also exhibited extensive communication with epithelial cells, such as *FTH1*/*FTL*-*SCARA5*, *EFNA1*-*EPHA3,* and *JAG1/NOTCH2* (Figure 7C). Some studies have shown that these signaling pathways can increase the expression of TF (*TWIST1* and *SOX4*) that control the fate of *POSTN*^+^ fibroblasts [35, 36], highlighting that epithelial cells may promote the development of *CFD*^+^ fibroblasts into *POSTN*^+^ fibroblasts. We detected high expression of *JAG1* in *COL17A1*^+^ epithelial cells and *NOTCH2* in *CFD*^+^ fibroblasts. We also found colocalization of *JAG1* and *NOTCH2* in the ST data, which indicated that *COL17A1*^+^ epithelial cells mediate the differentiation of *CFD*^+^ fibroblasts via *JAG1*/*NOTCH2* regulators (Figure S9A and C).

**Figure 7.**
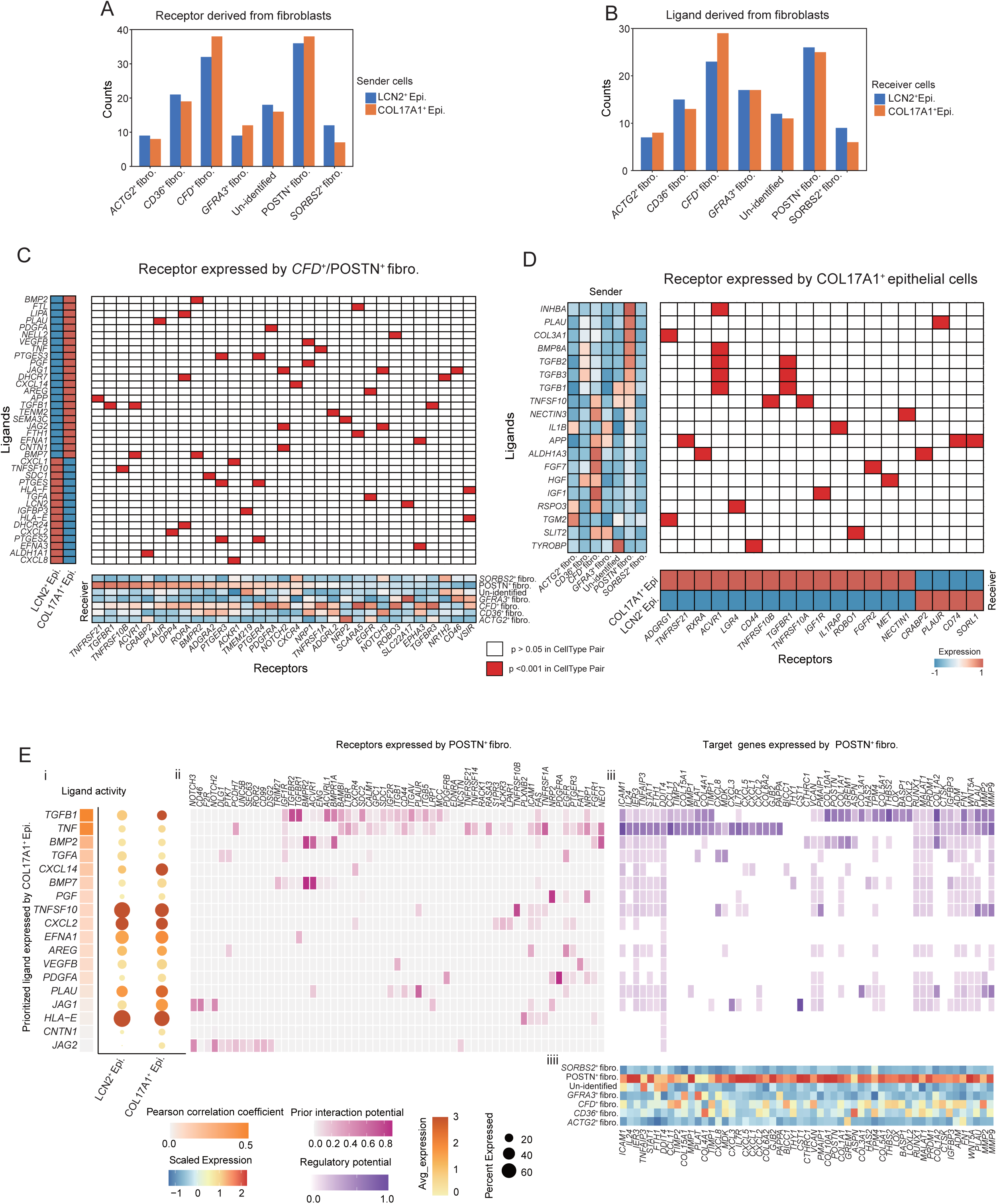
Ligand-receptor interaction between epithelial cells and fibroblasts. Bar plots of significant L-R pairs (*P* < 0.001, permutation test) from epithelial cell to fibroblast (**A**) or fibroblast to epithelial cell (**B**) in scRNA-seq data. **C.** Left: scRNA-seq average expression of epithelial-derived ligands modulating *POSTN*^+^ or *CFD*^+^ fibroblasts. Bottom: scRNA-seq average expression of corresponding receptors in *POSTN*^+^/*CFD*^+^ fibroblasts. Middle: heatmap of significant L-R pairs from *COL17A1*^+^ epithelial cells to *POSTN*^+^ or *CFD*^+^ fibroblasts. **D.** Left: heatmap showing scRNA-seq average expression of *POSTN*^+^ fibroblast-derived ligands modulating *COL17A1*^+^ epithelial cells. Bottom: heatmap of scRNA-seq average expression of corresponding receptors in *COL17A1*^+^ epithelial cells. Middle: heatmap of significant L-R pairs from *POSTN*^+^ fibroblast to *COL17A1*^+^ epithelial cell in scRNA-seq. **E.** (i) Dot plots showing the expression of top activity ligands in *COL17A1*^+^ epithelial cells from scRNA-seq; (ii) Heatmap of prior interaction potential scores of ligand-matched receptors in *POSTN*^+^ fibroblasts; (iii) Heatmap of regulatory potential scores of ligand matched target genes in *POSTN*^+^ fibroblasts; (iiii) Heatmap of scRNA-seq average expression of target genes in *POSTN*^+^ fibroblasts. L-R, ligand-receptor.

Fibroblasts serve as primary producers of ECM components and contribute to the formation of desmoplastic structures, which limit immune cell infiltration [37]. We found that *POSTN*^+^ fibroblast-specific genes were enriched in ECM-related pathways, which aligns with their role in the generation of desmoplastic structures (Figure S9B). CellPhoneDB and spatial proximity analysis revealed strong interactions between *COL17A1*^+^ epithelial cells and *POSTN*^+^ fibroblasts (Figure 7A and B). Therefore, we investigated whether *COL17A1*^+^ epithelial cells promote the ECM remodeling ability of *POSTN*^+^ fibroblasts. The results of the NicheNet analysis revealed that *COL17A1*^+^ epithelial cells presented high *TGFB1*, *TNF*, *BMP2*, and *TGFA* ligand activity (Figure 7E). These ligands interact with receptors on *POSTN*^+^ fibroblasts, inducing the expression of target genes encoding collagens or matrix metallopeptidases, such as *MMP1*, *MMP2, MMP9, COL11A1*, *COL15A1*, *COL4A1*, *COL6A2*, *COL5A1*, *COL10A1, COL1A1, COL1A2,* and *COL3A1,* in *POSTN*^+^ fibroblasts (Figure 7E). The results of the gene ontology (GO) enrichment analysis revealed that these target genes were enriched in terms of ECM organization and the collagen metabolic process, supporting our hypothesis (Figure S9D). In contrast, the top active TF in *POSTN*^+^ fibroblasts, including *HGF*, *PLAU*, *FGF7, INHBA, RSPO3, SLIT2, IL1B, IGF1, TGFB2,* and *TGFB3*, modulated the target genes in *COL17A1*^+^ epithelial cells, affecting pathways related to tumor progression, including epithelial cell proliferation, stem cell differentiation, and apoptotic signaling (Figure S9E and F). These findings indicate that the regulatory network between *POSTN*^+^ fibroblasts and *COL17A1*^+^ epithelial cells facilitates the progression of tumors and the formation of desmoplastic structures in a synergistic manner.

### *POSTN*^+^ fibroblasts producing *INHBA* promote tumor progression via the *ACVR1*/*SMAD2*/*TP63* axis

Cell–cell communication analysis revealed that *POSTN*^+^ fibroblasts play a key role in enhancing the malignant phenotype of tumor cells. To test this hypothesis, we conducted in *vivo* and in *vitro* experiments. Ligand-specific analysis revealed that among all possible ligands regulating tumor cells, *INHBA* had the highest expression specificity in *POSTN*^+^ fibroblasts (**Figure 8**A). Activin A (encoded by *INHBA*) is a secreted protein, and increasing activin A in tumor cells can promote tumor progression [38]. However, our data indicated that *INHBA* in the ESCC TME mainly originates from fibroblasts, suggesting that *INHBA* may be a key factor by which CAF promotes tumor progression (Figure 8B and C, Figure S10A). Additionally, *INHBA* was more highly expressed in tumor tissues than in normal tissues and was positively correlated with the master TF of *COL17A1*^+^ epithelial cells, *TP63* (Figure S10B and C). Therefore, we selected *INHBA* to conduct further experiments.

**Figure 8.**
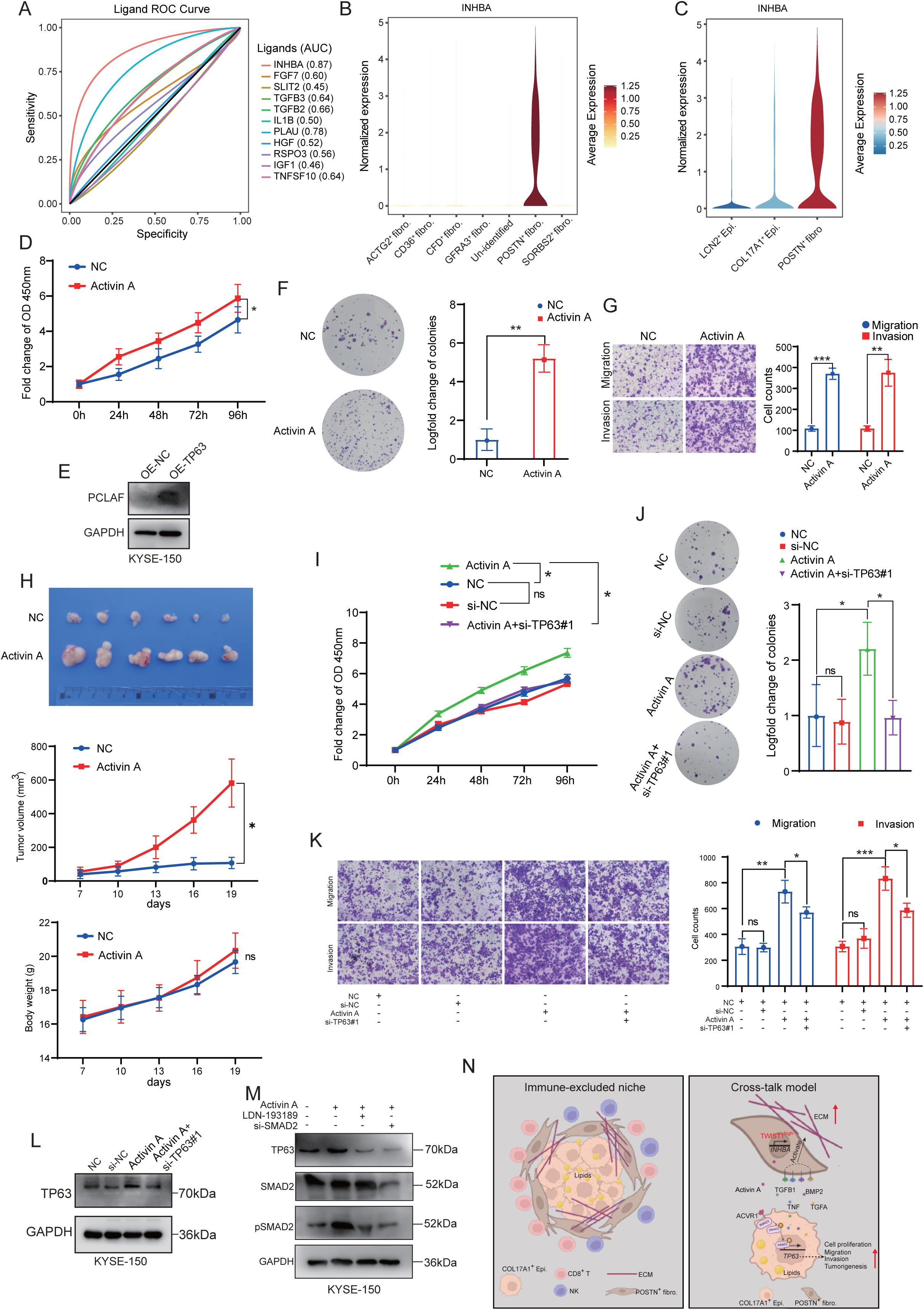
Validation of *INHBA* promoting tumor progression via the *ACVR1*/*SMAD2*/*TP63* axis. **A.** ROC curve analysis showing the specificity of top active ligands derived from *POSTN*^+^ fibroblasts. **B.** Violin plots showing mRNA expression levels of *INHBA* across seven fibroblast subpopulations. **C.** Violin plots showing mRNA expression levels of *INHBA* between epithelial subpopulations and *POSTN*^+^ fibroblasts. **D.** Cell viability was assessed at 0, 24, 48, 72, and 96 h using a CCK-8 assay in KYSE-150 cells treated with Activin A or negative control. **E.** WB analysis of *PCLAF* in KYSE-150 cells after treatment with Activin A or negative control. **F.** Quantification of colony formation efficiency after treatment of KYSE-150 cells with Activin A or negative control. **G.** Transwell experiments examining the migration and invasion abilities of KYSE-150 cells at 24 and 48 h after treatment with Activin A or negative control (n = 3). Scale bar = 100 μm. **H.** Representative images, changes of volume and body weight of subcutaneous tumor xenograft treated with negative control or Activin A in BALB/c nude mice (n=6). **I.** Cell viability was measured at 0, 24, 48, 72, and 96 h using a CCK-8 assay after treatment of KYSE-150 cells with Activin A or simultaneous treatment with si-*TP63* and Activin A. **J.** Quantification of colony formation efficiency after treatment of KYSE-150 cells with Activin A or simultaneous treatment with si-*TP63* and Activin A (n = 3). **K.** Transwell experiments examining the migration and invasion abilities of KYSE-150 cells at 24 and 48 h after treatment with Activin A or simultaneous treatment with si-*TP63* and Activin A (n = 3). Scale bar = 100 μm. **L.** WB analysis of *TP63* protein expression in KYSE-150 cells treated with Activin A or transfected with si-*TP63*. **M.** WB analysis of *TP63*, *SMAD2*, p-*SMAD2* protein expression in KYSE-150 cells treated with Activin A and transfected with si-*SMAD2* or *ACVR1* inhibitor LDN-193189. **N.** Immune-excluded niche model and cross talk model of epithelial and fibroblast cells. ns, *P* is not significant; *, *P* < 0.05; **, *P* < 0.01; ***, *P* < 0.001. Statistical significance was calculated by Student’s *t*-test, mean ± SD.

The results of the Cell Counting Kit-8 (CCK-8) assays revealed that treatment with the recombinant protein activin A significantly increased the viability of tumor cells (Figure 8D). Moreover, activin A also increased the expression of the cell proliferation marker *PCLAF* (Figure 8E). Clonogenic assays showed that activin A-treated tumor cells had enhanced colony-forming abilities (Figure 8F). Transwell assays also revealed that activin A treatment increased the migration and invasion capabilities of tumor cells, and xenograft experiments revealed that activin A enhanced the tumorigenic potential of tumor cells (Figure 8G and H). To determine whether activin A regulates the malignant phenotype of ESCC cells through *TP63*, we performed rescue experiments by simultaneously treating tumor cells with activin A and si-*TP63*, followed by monitoring changes in cell viability, colony formation, migration, and invasion ability. The results showed that knocking down *TP63* reversed the effects of activin A on promoting the malignant phenotype of tumor cells (Figure 8I–K). Moreover, simultaneous treatment of KYSE-150 cells with activin A and si-*TP63* reversed the ability of activin A to promote the expression of *TP63* (Figure 8L). Studies have shown that the *INHBA*-*ACVR1* pathway activates the *SMAD2* pathway, with *SMAD2* acting as a co-TF to promote gene expression [39]. Therefore, we determined whether *INHBA* promotes *TP63* expression via the *ACVR1*/*SMAD2* pathway. The results of gene manipulation experiments revealed that activin A promoted *TP63* expression and increased phosphorylated *SMAD2* levels (Figure 8M). Treatment with si-*SMAD2* or an *ACVR1* pathway inhibitor (LDN-193189) decreased *TP63* and phosphorylated *SMAD2* levels, indicating that activin A promoted *TP63* expression through the *ACVR1*/*SMAD2* pathway (Figure 8M). To summarize, *INHBA*, an essential oncogenic factor, is derived primarily from *POSTN*^+^ fibroblasts and promotes the malignant phenotype of tumor cells via the *ACVR1*/*SMAD2*/*TP63* axis in ESCC.

## Discussion

Although tumor immunotherapy has achieved remarkable curative outcomes in the treatment of various tumors, including ESCC [40], many patients who are administered immunotherapy develop treatment resistance or failure due to the heterogeneity of tumor cells and the TME [41]. Therefore, the key molecular events underlying the TME need to be determined to identify predictive biomarkers and optimize immunotherapeutic strategies. We performed scRNA-seq to construct a single-cell transcriptomic cell atlas within ESCC and integrated multimodal spatial profiling to assess tumor transcriptomic and metabolic reprogramming and the TME architecture. In this study, the mechanisms underlying the formation of the immunosuppressive microenvironment were elucidated. The heterogeneity and interactions between tumor cells and fibroblasts were systematically evaluated, emphasizing the key role of these interactions in tumor progression.

The TME can be classified into three distinct types: the inflamed type, the immune-excluded type, and the immune-desert type [42]. In ESCC, we found that CD8^+^ T cells coexpressed immune effector and immunosuppressive molecules, reflecting the immune-exhausted state in the TME [43]. NK cells are important antitumor cells with promising therapeutic prospects [44]. We found that *FGFBP2*^+^ NK cells were in a state of immune activation, as determined by their low expression of inhibitory molecules and favorable prognosis. However, the antitumor effect of *FGFBP2*^+^ NK cells may be limited by their low abundance in the TME. Therefore, our findings suggest that increasing the *FGFBP2*^+^ NK subpopulation in the TME may constitute an important antitumor strategy. Tregs play a crucial role in immune suppression [45], but their recruitment mechanism in the TME remains unclear. Our omics data revealed a pronounced chemotactic effect on Tregs in the TME via epithelial and stromal cells, including fibroblasts, mono/macrophages, and cDCs. Along with Tregs, Bregs are involved in the inhibitory microenvironment [46]. However, the functions and molecular characteristics of Bregs remain unclear because few relevant studies exist. In this study, B cells, particularly *JCHAIN*^+^ plasma cells, widely expressed the immunosuppressive molecules *TGFB1* and *IGHG4* compared to those in adjacent nontumor tissues. Moreover, some immunosuppressive molecules (*VISR, LGALS9,* and *TNFRSF4*) were highly expressed in B cells. These results indicate that B cells, like Tregs, participate in the formation of a suppressive immune microenvironment.

Cancer cells show differences in differentiation status, genomic variation, proliferation, and invasiveness between tumors and even within tumors [47]. Hence, the precise identification of some conserved key tumor cell subpopulations underlying tumor deterioration at the single-cell level may improve our understanding of tumorigenesis mechanisms and lead to the discovery of more universal therapeutic targets. We recapitulated two recurrent epithelial populations (*LCN2*^+^ epithelial cells and *COL17A1*^+^ epithelial cells) within ESCC via the comprehensive integration of single-cell spatial multiomics and an external single-cell dataset. *COL17A1*^+^ epithelial cells are characterized by the activation of the EMT pathway, strong proliferation ability, and poor prognosis. We also identified key TF determining the cell state of the two epithelial subpopulations. We found that TF *TP63* was significantly activated in *COL17A1*^+^ epithelial cells and considerably increased the expression of *COL17A1*^+^ epithelial cell marker genes, such as *KRT15*, *COL17A1*, and *EGFR*, indicating that *TP63* plays a key role in maintaining the state of *COL17A1*^+^ epithelial cells. Functional experiments revealed that *TP63* can significantly promote the malignant phenotype of esophageal cancer.

Some studies have shown that spatial information affects the transcriptomic reprogramming of tumors [48]. Consistent with this, the two epithelial subpopulations occupied distinct spatial niches, emphasizing the plasticity driven by spatial factors. Along with transcriptomic reprogramming, metabolic abnormalities are also hallmarks of the development of tumors [49]. Spatial metabolomics was recently used to elucidate the metabolic heterogeneity of the TME [50]. However, given the lack of metabolic markers for the identification of cell types, determining the metabolic features of specific tumor or stromal cell subpopulations is challenging. In this study, we proposed an approach to integrate spatial transcriptomic and metabolic data to illustrate the metabolic heterogeneity of the interested subpopulations, using single-cell transcriptomics as a reference. Using our customized workflow, metabolic heterogeneity was determined between *LCN2*^+^ epithelial cells and *COL17A1*^+^ epithelial cells. *COL17A1*^+^ epithelial cells are characterized by the accumulation of TG and abnormal PC metabolism. This study provides a framework for investigating metabolic alterations in refined subpopulations.

Cancer-associated fibroblasts play key roles in the solid tumors, influencing cancer progression [51]. Recent advancements in single-cell RNA sequencing have provided a comprehensive understanding of the complexity and heterogeneity of CAF subsets in various types of cancer [52]. In this study, we systemically deciphered the transcriptomic heterogeneity of CAF and identified a key protumorigenic fibroblast subpopulation, *POSTN^+^* fibroblasts, characterized by ECM remodeling related to desmoplastic structures. We also identified the key TF *TWIST1* in *POSTN*^+^ fibroblasts, which is closely associated with the function of *POSTN*^+^ fibroblasts. We identified an immune barrier composed of *POSTN*^+^ fibroblasts and *COL17A1*^+^ epithelial cells, with decreased infiltration of CD8^+^ T/NK cells and increased enrichment of lipids. Therefore, we proposed an immune exclusion model in which *POSTN*^+^ fibroblasts encapsulate *COL17A1*^+^epithelial cells, preventing immune cells from infiltrating the tumor core region. Additionally, the aberrant production of lipids by *COL17A1*^+^ epithelial cells further impairs the activity of immune cells (Figure 8N).

Given the unique niche formed by *COL17A1*^+^ epithelial cells and *POSTN*^+^ fibroblasts, we systematically mapped the mutual regulatory networks between the two cell types to elucidate the mechanisms underlying spatial niche formation. We found extensive cell communication between *COL17A1*^+^ epithelial cells and *POSTN*^+^ fibroblasts. We also found that ligands such as *TGFB1*, *TNF*, *BMP2*, and *TGFA* in *COL17A1*^+^ epithelial cells can enhance the matrix remodeling capabilities of *POSTN*^+^ fibroblasts, highlighting the role of *COL17A1*^+^ epithelial cells in the formation of this unique niche. In contrast, ligands such as *HGF*, *PLAU*, *FGF7*, *INHBA*, *RSPO3*, *SLIT2*, *IL1B*, *IGF1*, *TGFB2*, and *TGFB3* in *POSTN*^+^ fibroblasts can promote the proliferation and differentiation of *COL17A1*^+^ epithelial cells. Our in *vitro* and in *vivo* experiments demonstrated that the ligand *INHBA*, derived from *POSTN*^+^ fibroblasts, can promote the expression of the key TF *TP63* in *COL17A1*^+^ epithelial cells via the *ACVR1*/*SMAD2* pathway, suggesting that *POSTN*^+^ fibroblasts can facilitate the conversion of tumor cells into the *COL17A1*^+^ epithelial phenotype. Therefore, based on the above results, we proposed an interaction model between *POSTN*^+^ fibroblasts and *COL17A1*^+^ epithelial cells (Figure 8N).

## Conclusions

To summarize, this study provides a comprehensive single-cell spatial multimodal resource for ESCC. We systematically illustrated the suppressive immune microenvironment in ESCC. We also investigated the heterogeneity of tumor cells and fibroblasts and elucidated the role of interactions between key tumor cells and fibroblast subpopulations in the formation of the immune barrier and the promotion of tumor progression.

## Materials and methods

### Collection of clinical human patient samples

In this study, five ESCC tumor tissues and three patient-matched adjacent tissues were collected from the First Affiliated Hospital of Zhengzhou University. The patients were 65–69 years old. After detaching the tissue, it was immediately transferred to the laboratory for further processing. The inclusion criteria for patients were patients who were diagnosed with ESCC by a pathologist and had no other underlying diseases. The study was approved by the Ethics Committee of Scientific Research and Clinical Trials of the First Affiliated Hospital of Zhengzhou University (Approval 2022-KY-1544–001). All participants provided signed informed consent.

### Tissue dissociation and single-cell library construction

Fresh tissues were immediately preserved in a tissue preservation solution (Singleron) on ice. Tissues were dissociated following the manufacturer’s protocol (Singleron). The scRNA-seq library was constructed using GEXSCOPE® single-cell RNA library kits (Singleron) [53].

### Spatial transcriptomics

#### Tissue optimization

Tissue optimization was performed for ESCC tissues following the manufacturer’s recommendations (10× Genomics). Briefly, eight optimal cutting temperature compound (OCT) embedded cryosections (10 μm thick) were placed on spatially optimized slides and treated with permeabilization enzymes (10× Genomics) for different durations to determine the optimal penetration time. After determining the optimal conditions, the tissue sections were cut into 10-μm-thick sections and mounted onto spatial gene expression slides.

#### Fixation, staining, and imaging

Slides with sections were fixed in methanol for 20 min after incubation at 37 °C for 1 min. Before staining, the slides were incubated in isopropyl alcohol (l9516, Sigma) for 1 min. Then, the sections were incubated in Mayer’s hematoxylin (MHS16, Sigma) for 7 min and washed 15 times in ultrapure water. Next, the sections were incubated in bluing buffer (Dako) for 1 min, and eosin (HT110216, Sigma) diluted 1:10 in Tris-base (0.45 M Tris, 0.5 M acetic acid, pH 6.0) was added for 1 min. After air-drying, the slides were imaged at 20 × magnification using a slide scanning platform.

#### Permeabilization

The slides with tissue sections were inserted into cassettes to establish individual reaction chambers for each section. For permeabilization, the sections were incubated at 37 °C for 30 min with permeabilization enzymes. After incubation, the permeabilization enzyme was aspirated, and the wells were rinsed with 0.1× saline sodium citrate buffer.

### Spatial transcriptomic library preparation and sequencing

After permeabilization, cDNA synthesis and amplification (17 cycles) were performed using the Spatial Visium Reagent Kit (10× Genomics). The cDNA libraries were subsequently prepared using a spatial cDNA library kit (10× Genomics). The cDNA libraries were sequenced on the Illumina NovaSeq Xplus platform using an S4 Reagent Kit (Illumina).

### Spatial metabolomics

#### Sample preparation

The OCT-embedded tissues were sliced at a thickness of 14 _μ_m using a Leica CM3050S cryostat (Leica Microsystems GmbH, Wetzlar, Germany) at –20 °C. The sections were subsequently mounted on electrically conductive slides coated with indium tin oxide (ITO) and then dried in a vacuum desiccator for 30 min.

#### Matrix coating and mass spectrometry imaging

Dehydrated samples were treated with an HTX TM sprayer (Bruker Daltonics, Germany), and 15 mg/mL 2,5-dihydroxybenzoic acid solution mixed in 90% acetonitrile was applied. Next, a prototype Bruker timsTOF flex mass spectrometry imaging system (Bruker Daltonics, Bremen, Germany), equipped with a 10 kHz smartbeam 3D laser, was used for MALDI timsTOF MSI analysis. The laser power was consistently maintained at 80% throughout the process. Positive ion mode was selected for obtaining mass spectra within a mass range of 50–1300 Da. The tissue images were resolved at a spatial resolution of 50 _μ_m, with each spectrum compiled from 400 laser shots. Normalization was performed using the root mean square method on the MALDI mass spectra, and the signal intensity displayed in each image represented the normalized values. To confirm the structural identification of the detected metabolites, MS/MS fragmentation was conducted using the timsTOF flex MS system in the MS/MS mode.

### Immunofluorescence assay

The thickness of the paraffin sections was set at 5 _μ_m. For immunofluorescence staining, the sections were treated with xylene for deparaffinization (2 × 15 min) and subsequently rehydrated with a gradient of alcohol (100%, 95%, 80%, and 70%), followed by distilled water; all procedures were performed at room temperature. The sections were subjected to antigen retrieval at 95 °C for 30 min with sodium citrate buffer (0.01 M, pH 6.0). Then, the samples were sequentially blocked for 25 min and 30 min at 25 °C in 3% hydrogen peroxide solution and bovine serum albumin (BSA) buffer (0.3% Triton X-100, 1% BSA) and then rinsed with phosphate buffered saline (PBS) (3 × 5 min). The samples were subsequently treated for 10 h at 4 °C with primary antibodies and then washed three times with PBS (3 × 15 min). The samples were treated for 50 min with an horseradish peroxidase (HRP) labeled secondary antibody (Servicebio), followed by washing three times with PBS (3 × 5 min). The samples were treated for 10 min with tyramide (Servicebio) and then washed three times with tris buffered saline tween (3 × 5 min). Finally, the cell nuclei were stained with DAPI (G1012, Servicebio) for visualization, and fluorescence imaging was performed. The primary antibodies used included rabbit anti-CD16 (1:3000, GB113963, Servicebio), mouse anti-CD8α (1:5000, GB12068, Servicebio), rabbit anti-Periostin (1:500, ab215199, Abcam), rabbit anti-COL17A1 (1:100, ab184996, Abcam), and rabbit anti-Lipocalin-2 (1:3000, #44058, CST) antibodies. The secondary antibodies used included goat HRP-labeled anti-IgG (1:500, GB23301 and GB23303, Servicebio). Goat anti-rabbit IgG (1:2000, A-11008, Thermo Fisher) was used. Tyramide included iF488-tyramide (1:500, G1231, Servicebio), iF555-tyramide (1:500, G1233, Servicebio), and iF647-tyramide (1:500, G1232, Servicebio).

### Cell culture

The KYSE-150 and BJ cell (human fibroblast) lines were purchased from the Cell Bank of the Chinese Academy of Sciences (Shanghai, China). KYSE-150 cells were cultured in RPMI-1640 medium, while BJ cells were cultured in DMEM (Gibco), both supplemented with 10% fetal bovine serum (FBS) (Invigentech). The cells were incubated at 37 °C in a humidified atmosphere with 5% CO_2_.

### Cell proliferation, colony formation, and cell migration and invasion assays

Cell proliferation was assessed at predetermined intervals using a CCK-8 (K1018, APExBIO), following the manufacturer’s instructions. For the colony formation assay, cells were seeded in six-well plates at a density of 1000 cells per well in RPMI-1640 medium supplemented with 10% FBS (Gibco). The cells were incubated at 37 °C in a humidified atmosphere containing 5% CO_2_ for 14 days. The colonies were subsequently fixed with 4% paraformaldehyde and stained with crystal violet, and the visible colonies were counted. For the migration assay, cells (5 × 10^4^ cells per well) were suspended in 200 μL of FBS-free RPMI-1640 medium and added to a Transwell insert (Corning). The lower compartment of each well was filled with 500 μL of RPMI-1640 containing 10% FBS. For the invasion assay, Transwell chambers were precoated with 4% Matrigel (BioCoat) diluted in RPMI-1640 and incubated at 37 °C for 1 h. The cells were then seeded in the upper chamber as described in the migration assay. After 48 or 72 h, the Transwell inserts were collected and stained with 0.5% crystal violet. Images of the stained cells on the lower surface of the inserts were captured, and the number of cells in three random fields was quantified under a microscope.

### Western blotting

Total proteins were extracted using RIPA lysis buffer (P0013B, Beyotime, China), and their concentrations were quantified by conducting the bicinchoninic acid (BCA) protein assay. The following primary antibodies were used in this study: anti-p63 (Proteintech, 12143-1-AP), anti-COL17A1 (Abcam, ab184996), anti-Cytokeratin 15 (Proteintech, 10137-1-AP), anti-EGFR (Proteintech, 18986-1-AP), anti-GAPDH (Proteintech, 10494-1-AP), anti-SMAD2 (Proteintech, 12570-1-AP), anti-pSMAD2 (Abcam, 280888), anti-Activin A (Proteintech, 60352-1-lg), anti-TWIST1 (Proteintech, 25461-1-AP), and anti-Periostin (Abcam, ab215199).

### RNA interference

The siRNAs were prepared in Opti-MEM (Gibco, USA) and introduced into the cell lines using Lipofectamine 3000 (Invitrogen) following the manufacturer’s protocol.

### *In vivo* tumorigenesis and metastasis assays

Null/null athymic BALB/c female mice (5–6 weeks old) were purchased from Beijing Vital River Laboratory Animal Technology Co., Ltd. (Beijing, China), and all procedures were performed according to the Animal Research Reporting of In Vivo Experiments (ARRIVE) guidelines. For xenograft implantation, KYSE-150 cells (5 × 10^6^ cells per mouse) were suspended in PBS and subcutaneously injected into the right flank of each mouse. After rearing for three days, the mice were administered 4 mg/kg recombinant activin A (Thermo, P08476) in 0.9% NaCl via intraperitoneal (i.p.) injection every two days. Tumor volumes were measured every two days using calipers, with calculations based on the formula: V = (L × W^2^) / 2, where L represents the longest diameter and W the perpendicular shorter diameter. After feeding for 20 days, the mice were euthanized, and the tumors were harvested for further analysis.

### Quantitative real-time polymerase chain reaction

Total RNA was isolated using TRIzol reagent (Invitrogen) and subsequently reverse-transcribed into cDNA using the PrimeScript RT Reagent Kit with gDNA Eraser (Takara). Quantitative real-time polymerase chain reaction (qPCR) was conducted using TB Green® Premix Ex Taq™ II (Tli RNaseH Plus) (Takara) following the manufacturer’s instructions. Next, 18S was used as an internal control, and gene expression levels were quantified using the 2^−ΔΔCt^ method.

### Construction of stably overexpressing cell lines

The *TP63* and *TWIST1* lentiviruses were constructed by GENECHEM (Shanghai, China). Then, *TP63* and *TWIST1* lentiviruses were transfected into KYSE-150 and BJ cells using HitransG P solution (GENECHEM) for 24 h. Positive cells were selected with puromycin (2 µg/mL, Solarbio, P8230) and subsequently verified by conducting Western blotting and qPCR assays.

### Raw sequence data processing

Human reference genome data were obtained from the 10× Genomics website (https://cf.10xgenomics.com/supp/cell-exp/refdata-gex-GRCh38-2020-A.tar.gz). CellRanger software (10× Genomics) and Spaceranger software (10× Genomics) were used to preprocess the raw sequencing data to generate UMI counts for the scRNA-seq and spatial transcriptomics data, respectively.

### Processing of scRNA-seq matrix data

Gene expression data were loaded into the R platform for analysis using the Seurat software [54]. Low-quality cells with fewer than 200 detected genes and more than 20% mitochondrial genes were removed before further analysis. Genes whose expression was under three cells were filtered out. The gene-barcode count matrices were normalized using the ‘SCTransform’ function from Seurat. To decrease dimensionality, the top 3000 genes with high variability were identified using the ‘FindVariableGenes’ function. Principal component analysis (PCA) was performed for dimensionality reduction. Then, all matrices were integrated into a combined dataset, and the batch effects from the samples were removed using the robust principal component analysis (RPCA) algorithm. UMAP analysis was performed using the ‘RunUMAP’ function with the top 30 principal components (PCs) as inputs. Cluster analysis for total cells was performed using the Louvain algorithm (resolution = 0.5).

### Spatial transcriptomics clustering analysis

Spatial gene expression and image data were analyzed using the R package Seurat. Spots not covered by the tissue section were removed from downstream analysis. The ‘SCTransform’ function was used to normalize the raw spot-barcode matrices. Dimensionality reduction was performed using the PCA algorithm for each spatial sample. Clustering analysis was also performed using the Louvain algorithm (resolution = 0.5) based on UMAP plots. Spot clusters were visualized in UMAP plots or spatial spots over H&E images. Factor analysis was conducted to determine the abundance of distinct cell populations in each spot using the ‘FindTransferAnchors’ and ‘TransferData’ modules in the Seurat software. The ‘SpatialFeaturePlot’ function in Seurat was used to visualize the enrichment of distinct cell types.

### Copy number variation analysis

Using the pyinfercnv software, CNV analysis was performed on epithelial cells. Briefly, the raw single-cell matrix was used as the input file, and 1,000 immune cells were randomly selected as the reference. The standard pyinfercnv pipeline was run to calculate CNV values.

### Identification of differentially expressed genes

Cell-specific marker genes were determined using the ‘Findallmarker’ function in the Seurat software for each cell population. For a given cell type, the genes expressed in more than 10% of the cells with a log_2_-fold change > 0.25 were regarded as marker genes. Using the ‘Findmarker’ function in the Seurat software, spatial region-restricted genes were identified by comparing each compartment of interest in the tissue sections. Genes with log_2_-fold change > 0.5 and adjusted *P* < 0.05 were regarded as statistically significant spatially variable genes.

### Cell pseudotime analyses

Two methods were used for cell trajectory inference. (1) The cell development trajectory was inferred using the R package Monocle2. First, the single-cell UMI count matrices were normalized using the ‘estimateSizeFactors’ and ‘estimateDispersions’ modules in Monocle2. Genes with a mean value > 0.1 were retained for dimensionality reduction and generation of the cell trajectory. Finally, the cell pseudotime trajectory was established using the ‘DDRTree’ method in Monocle2. (2) The scRNA-seq data were preprocessed to obtain spliced and unspliced RNA counts for each gene in each cell, followed by normalization and filtering of low-quality cells and genes. Using the velocyto R package, RNA velocity vectors were estimated by computing the ratio of spliced to unspliced mRNA for each gene, and these vectors were projected onto a low-dimensional embedding to visualize inferred cell trajectories.

### GO and pathway enrichment analysis

GO and pathway enrichment analyses were performed using the R package clusterProfiler [55]. Gene identifiers were mapped using the annotation Dbi R package org.Hs.eg.db. All expressed genes were included as backgrounds. GO items with *P* < 0.05 were considered to be statistically significant. The results were visualized using the ggplot2 R package.

### Cell interaction analysis

L-R pairs were scanned using the CellPhoneDB [56] tool. First, we computed the average mRNA expression of ligand-receptor pairs across distinct cell–cell pairs in single-cell sequencing data. Only genes expressed in more than 10% of the cells per cell were considered available ligands or receptors. Then, CellPhoneDB was run to determine the available L-R pairs for each cell-cell pair, with default parameters and normalized count matrices as inputs. For statistically significant L-R pairs, the threshold was set at an average expression level > 0.2 and *P* < 0.001.

### Transcription factor regulon analysis

The SENIC [32] tool was used to determine the active transcription regulon in cell types of interest. A total of 2025 unique TFs were obtained by integrating resources from four TF databases, including JASPAR (http://jaspar.genereg.net/)[57], AnimalTFDB (https://guolab.wchscu.cn/AnimalTFDB4/) [58], hTFtarget (http://bioinfo.life.hust.edu.cn/hTFtarget#!/) [59] and TRRUST (https://www.grnpedia.org/trrust/) [60]. The hg38 human reference genome was obtained from a website (https://resources.aertslab.org/cistarget/databases/homo_sapiens/hg38/refseq_r80/mc9 nr/gene_based/). First, we calculated the correlation between TF-target pairs to build a coexpression module using the GENIE3 algorithm in normalized scRNA-seq data. The TF regulons were subsequently determined according to the hg38 human genome reference data. Finally, the R package AUCell was used to calculate area under the curve (AUC) scores representing the transcriptional activity of the TF regulons. For cell-type-specific TF regulons, we used the Shannon entropy algorithm to determine the top-specific TF regulons for each cell cluster based on the AUC scores.

### Cell preference distribution analyses

To evaluate the differences in the distributions of fibroblast or epithelial cell subpopulations in cancer cells and adjacent tissues, the Ro/e value was calculated to represent the degree of tissue preference distribution. For a given cell subpopulation i and its corresponding tissue j, a 2 × 2 contingency table was constructed. This table included the number of cells i in tissue j and other tissues. The number of non-i cells in tissue j and in other tissues was also counted. A Fisher test was subsequently performed on the contingency table to obtain the Ro/e value and the corresponding p-value. A Ro/e value > 1 indicates that cell subpopulation i is more likely to be distributed in tissue j, whereas a lower Ro/e value < 1 indicates that cell subpopulation i is less likely to be distributed in tissue j.

### Alignment of metabolic and spatial transcriptomics data

To integrate spatial transcriptomic and metabolomic data, we developed an image-based integration method. Briefly, we employed an image registration pipeline involving affine alignment and nonlinear transformation to spatially co-register transcriptomic and metabolomic images. We used the simple ITK framework (Python software used for medical image analysis) for image preprocessing and registration (https://simpleitk.org/). First, the ST H&E and SM images were converted to grayscale images and resampled to the same size. Then, we aligned the centers of the two images and registered the images via affine transformation. A mutual information algorithm was used to evaluate similarity. The optimizer was established using the gradient descent algorithm. Through this approach, we obtained the transformation parameters of the registration. Finally, the pixel coordinates of the ST and SM images were converted to the same coordinate system. Because of the distinct resolution between the ST and SM data, it was necessary to integrate the ST spots and SM pixels. For a given ST spot *S* = 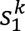, i = 1, …k, with n surrounding metabolic pixels *M*= 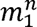, i = 1, …n. The distance between S and M is *d*_|*S-M*|_ < 100 µm. We defined the metabolic intensities of spot S *M_s_* using the following formula:

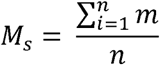

### Identification of differentially abundant metabolites

The ‘FindAllMarkers’ function in Seurat was used to compute differentially abundant metabolites in the four spatial niches. To identify differentially abundant metabolites between *LCN2*^+^ and *COL17A1*^+^ epithelial cells, the proportion of epithelial cell subpopulations in each spot in the epithelial region was first calculated using the deconvolution algorithm in Seurat. Next, the average proportion of each epithelial cell subpopulation in each spatial cluster in the epithelial region was determined. Spatial seurat clusters with high average proportions were classified as corresponding epithelial cell subpopulation spatial clusters. Finally, the ‘FindAllMarkers’ function in Seurat was used to calculate the differentially abundant metabolites of the epithelial cell subpopulations. Log_2_-fold change > 1 and *P* < 0.05 were considered statistically significant.

### Cellular neighborhood detection

The method used by Dalia et al. was used to detect the neighborhood of a cell in spatial data [61]. We defined the ‘neighborhood score’ as the fraction of surrounding spots containing that cell type, directly measuring the cell type composition in adjacent spots. The neighborhood of a spot was defined as spots with a distance less than or equal to 1 (including the spot), resulting in sets of no more than seven neighbors per spot. The neighborhood cell type fraction was calculated from the binarized cell type annotations of this set. Briefly, we first calculated the neighborhood score for each fibroblast subgroup in the barrier region. We subsequently used the deconvolution algorithm to calculate the proportion of *LCN2*^+^ and *COL17A1*^+^ epithelial cells in each spot of the barrier region. Finally, we counted the number of neighborhood spots, defined as the proportion of *LCN2*^+^ or *COL17A1*^+^ epithelial cells, and the neighborhood score of each fibroblast subgroup was greater than 0.1 to evaluate the cellular neighborhood.

### Analysis of TCGA expression and survival curves

The ESCC bulk transcriptomic data with clinical information, including 95 tumor samples and 13 normal samples, were downloaded from the cancer genome atlas (TCGA) database (https://www.cancer.gov/ccg/research/genome-sequencing/tcga) using the tool TCGAbiolinks [62]. The read count matrices were normalized using the transcripts per million (TPM) method. The TPM data were subsequently used to detect the expression of genes of interest. For survival analysis, 75 tumor samples remained after the samples without clinical information were filtered out. Survival curves were drawn using the R package survival. A Kaplan–Meier survival curve was generated using the ‘survfit’ function in survival software.

## Ethical statement

The animal experiments were approved by the Ethics Committee of Scientific Research and Clinical Trial of the First Affiliated Hospital of Zhengzhou University (Approval number: ZZU-LAC20240531). The sample collection was approved by the Ethics Committee of Scientific Research and Clinical Trial of the First Affiliated Hospital of Zhengzhou University (Approval number: 2022-KY-1544-001).

## Data availability

The raw data reported in this paper have been deposited in the Genome Sequence Archive at the National Genomics Data Center, Beijing Institute of Genomics, Chinese Academy of Sciences/China National Center for Bioinformation (GSA-Human: HRA007162), and are publicly accessible at https://ngdc.cncb.ac.cn/gsa-human. The main code for this study is stored in https://github.com/biosyy/single-cell-and-spatial-transcriptomic.

## CRediT author statement

**Yong Shi:** Writing - original draft, Validation, Methodology, Formal analysis, Resources. **Ke An:** Software, Formal analysis. **Yu Qi:** Resources, Investigation. **Xinhan Zhang:** Validation. **Yueqin Wang:** Validation, Investigation. **Xuran Zhang:** Resources, Data Curation. **Shaoxuan Zhou:** Writing - review & editing. **Ouwen Li:** Writing - review & editing. **Yanan Song:** Writing - review & editing. **Jiayi Zhou:** Writing - review & editing. **Yue Du:** Writing - review & editing. **Mingyang Hou:** Writing - review & editing. **Yun-Gui Yang:** Conceptualization, Supervision. **Quancheng Kan:** Conceptualization, Supervision. **Xin Tian:** Funding acquisition. Conceptualization, Supervision. All authors have read and approved the final manuscript.

## Competing interests

The authors declare no conflicts of interest.

## Acknowledgements

This work was supported by Funding for Scientific Research and Innovation Team of The First Affiliated Hospital of Zhengzhou University (ZYCXTD2023010), the Science and Technology Innovation Leading Talent Program of Henan Province (No. 254000510028), and the Special Fund for Young and Middle School Leaders of Henan Health Commission (HNSWJW-2020017). This study was supported by the General Program of the National Natural Science Foundation of China (32170594).

## Supplementary material

**Table S1 Baseline information of clinical pathological parameters in five samples with ESCC**

**Figure S1.**
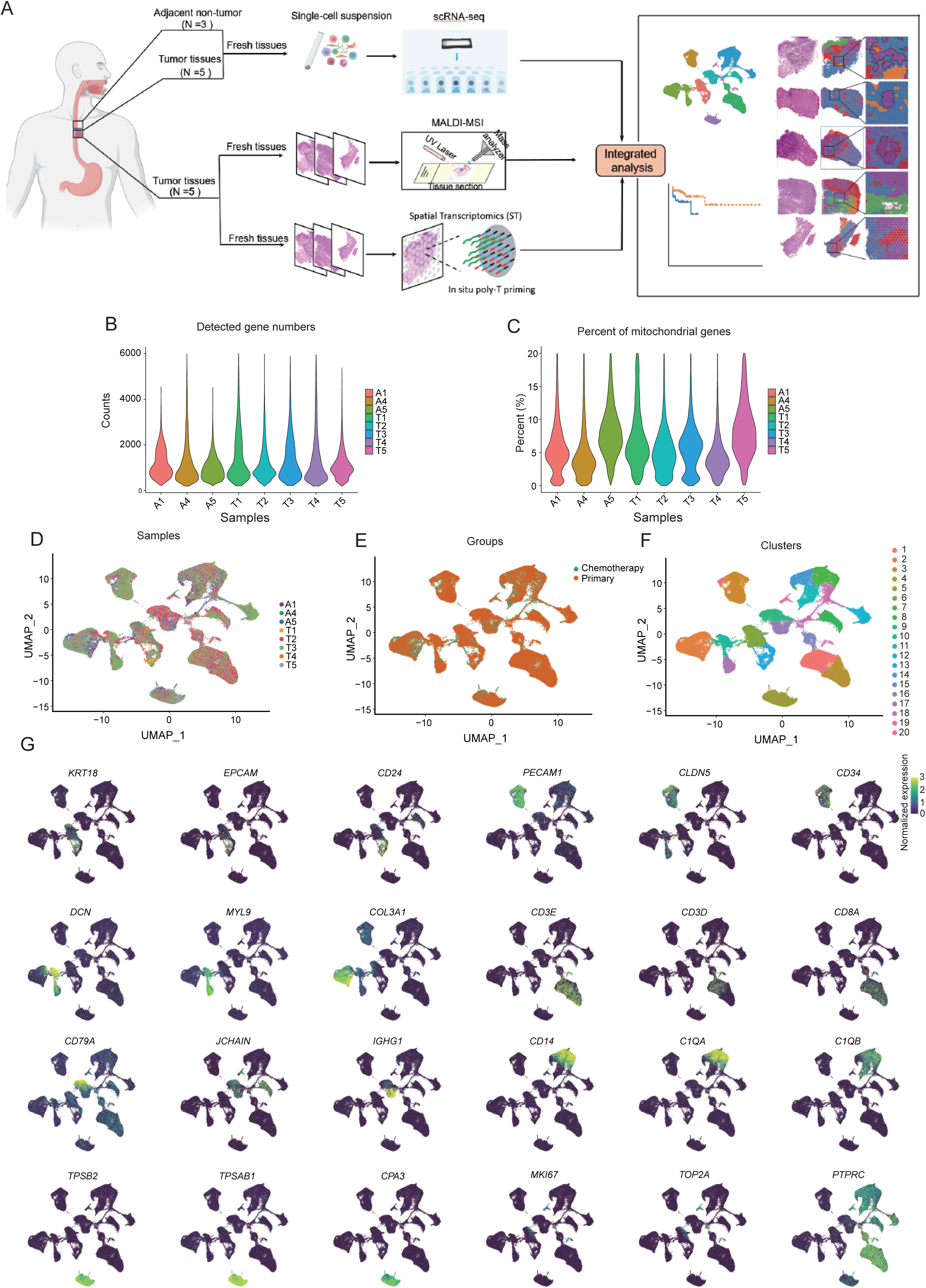
Single cell clustering and cell proportion analysis. **A.** Workflow of multiomics integration analyses for scRNA-seq, ST and SM. **B.** Violin plots showing average counts of detected genes in each ESCC sample. **C.** Violin plots showing the percent of mitochondrial genes across all samples after quality control. **D.** UMAP plots of total cells labeled by samples. **E.** UMAP plots of total cells labeled by groups. **F.** UMAP plots of total cells labeled by seurat clusters. **G.** Feature plots of marker genes for eight major cell types.

**Figure S2.**
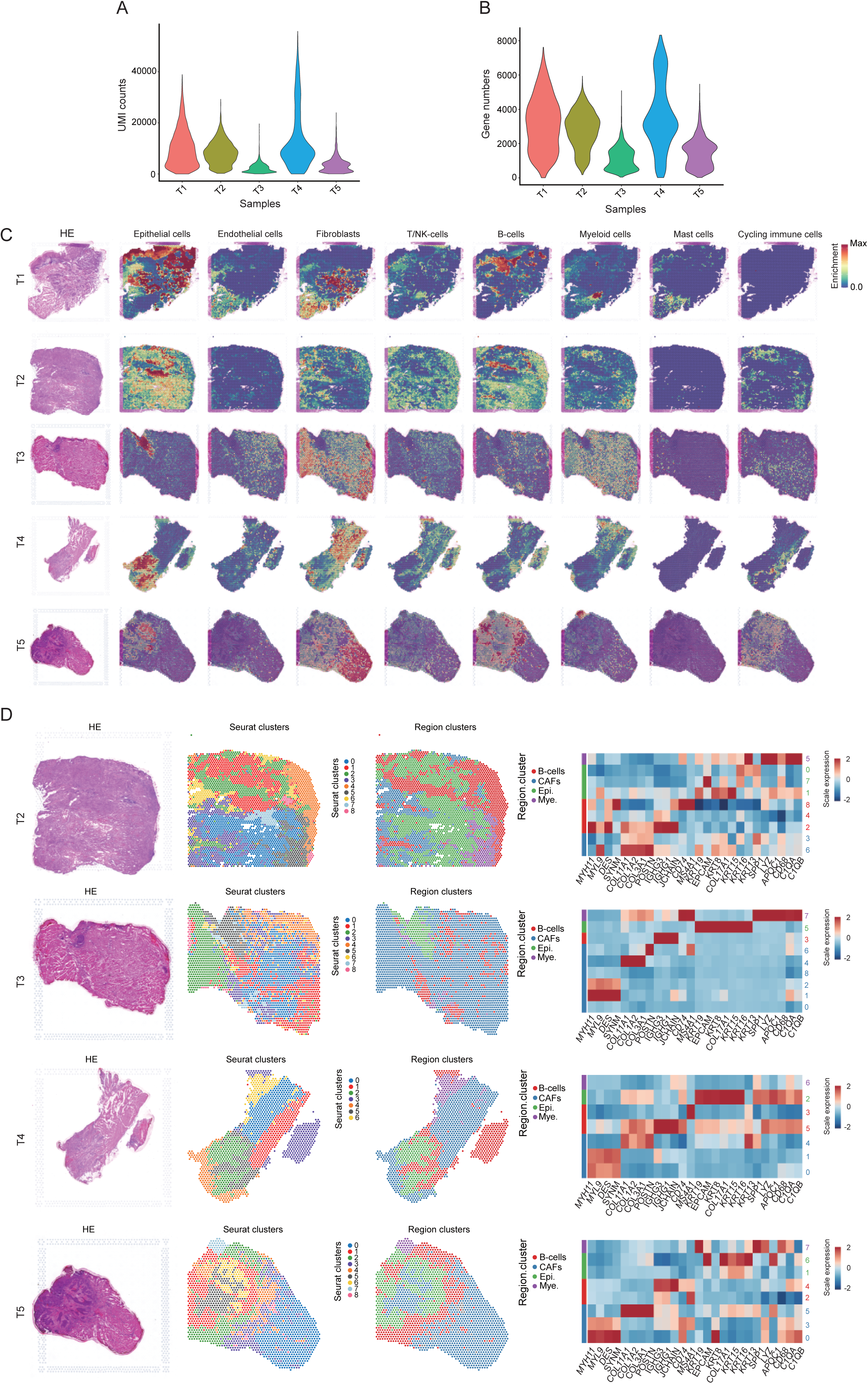
Integration of single-cell and spatial transcriptomic data. Violin plots showing the average UMI counts (**A**) and detected genes (**B**) across five sections in ST data. **C.** Spatial feature plots of enrichment scores for eight ESCC cell populations across five tumor samples. **D.** Left: H&E staining of tissue sections, unbiased clustering of ST spots and four spatial niche clusters. Right: heatmap showing the expression of marker genes for four spatial niches. UMI, unique molecular identifier.

**Figure S3.**
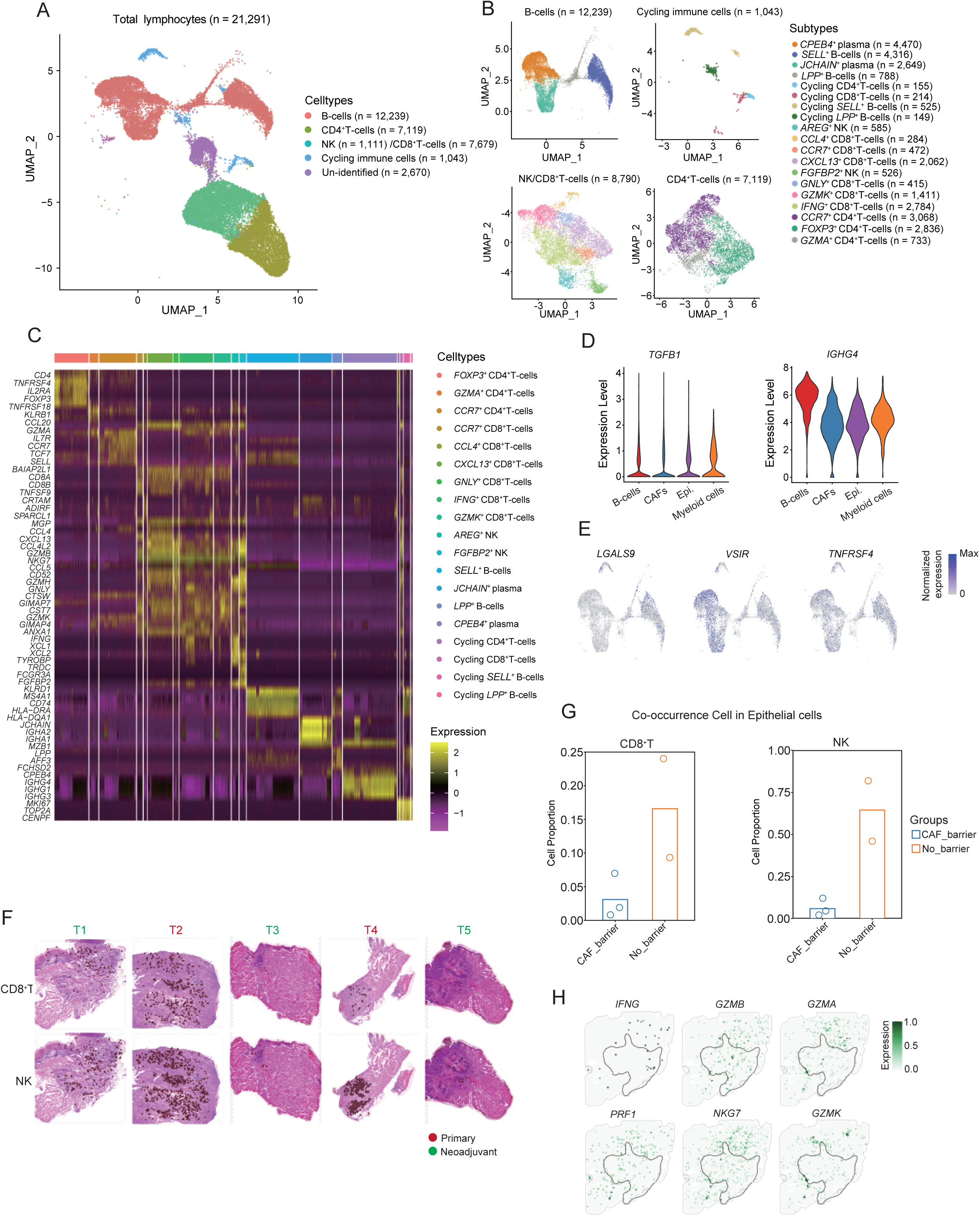
The features of tumor microenvironment in ESCC. **A.** UMAP plots of major lymphocyte populations. **B.** UMAP plots of subpopulations for B cells, cycling immune cells, CD8^+^ T cells, CD4^+^ T cells. **C.** Heatmap showing the mRNA expression of maker genes for all lymphocyte subpopulations. **D.** Violin plots showing the mRNA expression of *TGFB1* and *IGHG4* across four spatial niches in ST data. **E.** Feature plots of *LGALS9*, *VSIR*, *TNFRSF4* for B cells in scRNA-seq data. **F.** H&E staining with co-occurrence spots of CD8^+^ T/NK and epithelial cells. **G.** Box plots showing the proportion of co-occurrence spots of CD8^+^ T/NK and epithelial cells in total tumor spots between CAF barrier and no barrier groups. **H**. Spatial feature plots of selected immune effector genes including *IFNG*, *GZMB*, *GZMA*, *PRF1*, *NKG7*, *GZMK* in tumor sample 1.

**Figure S4.**
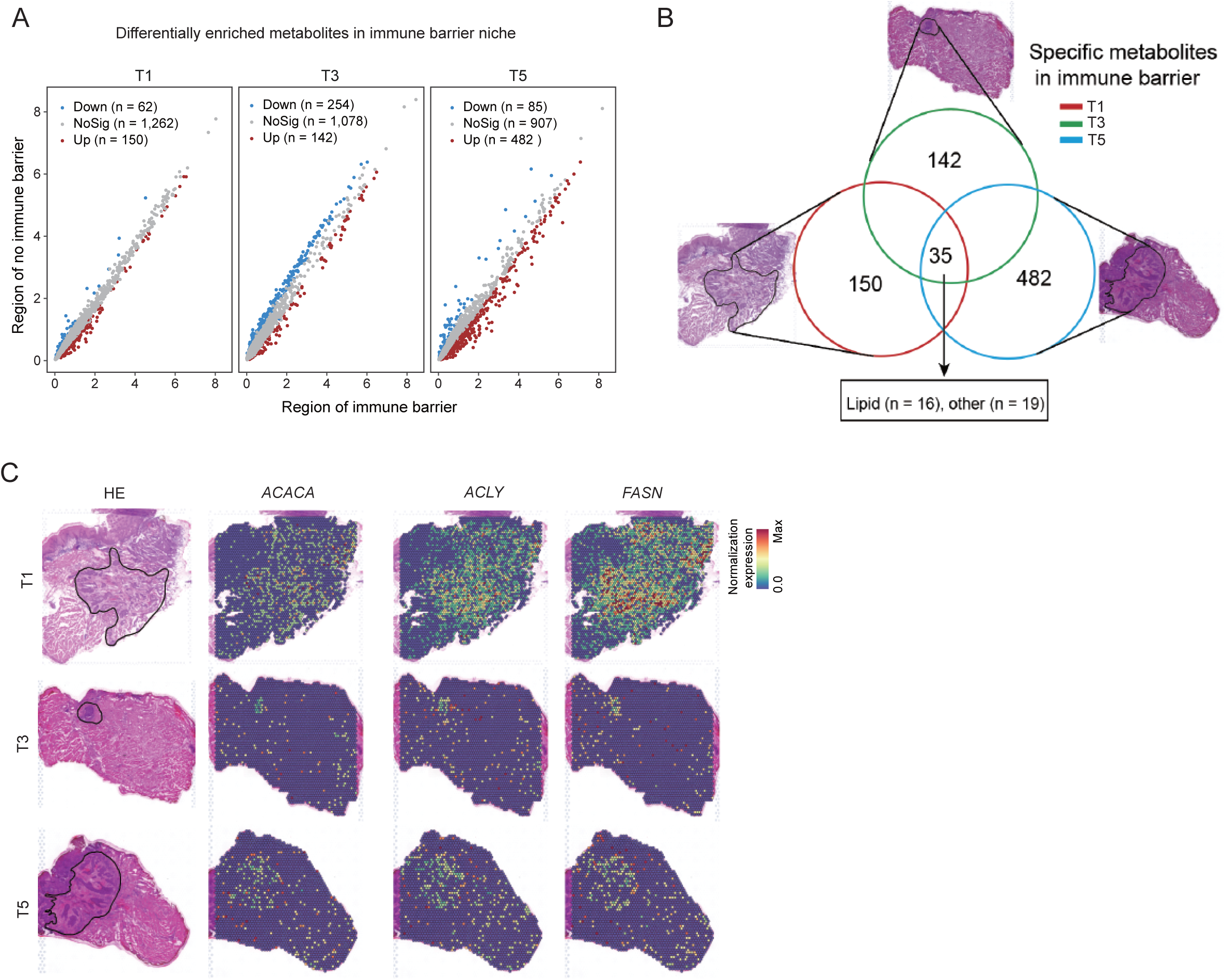
Identification of metabolic features in the immune barrier region. **A.** Volcano plots of differential metabolites (|log_2_-fold change| > 1 and *P* < 0.05) between the CAF barrier and non-barrier regions in tumor sample 1, 3, and 5. **B.** Venn diagrams illustrating the intersection of highly enriched metabolites in the immune barrier region across tumor sample 1, 3, and 5. **C.** Spatial feature plots displaying the spatial distribution of lipid synthesis-related genes *ACACA*, *ACLY*, and *FASN* in tumor sample 1, 3, and 5.

**Figure S5.**
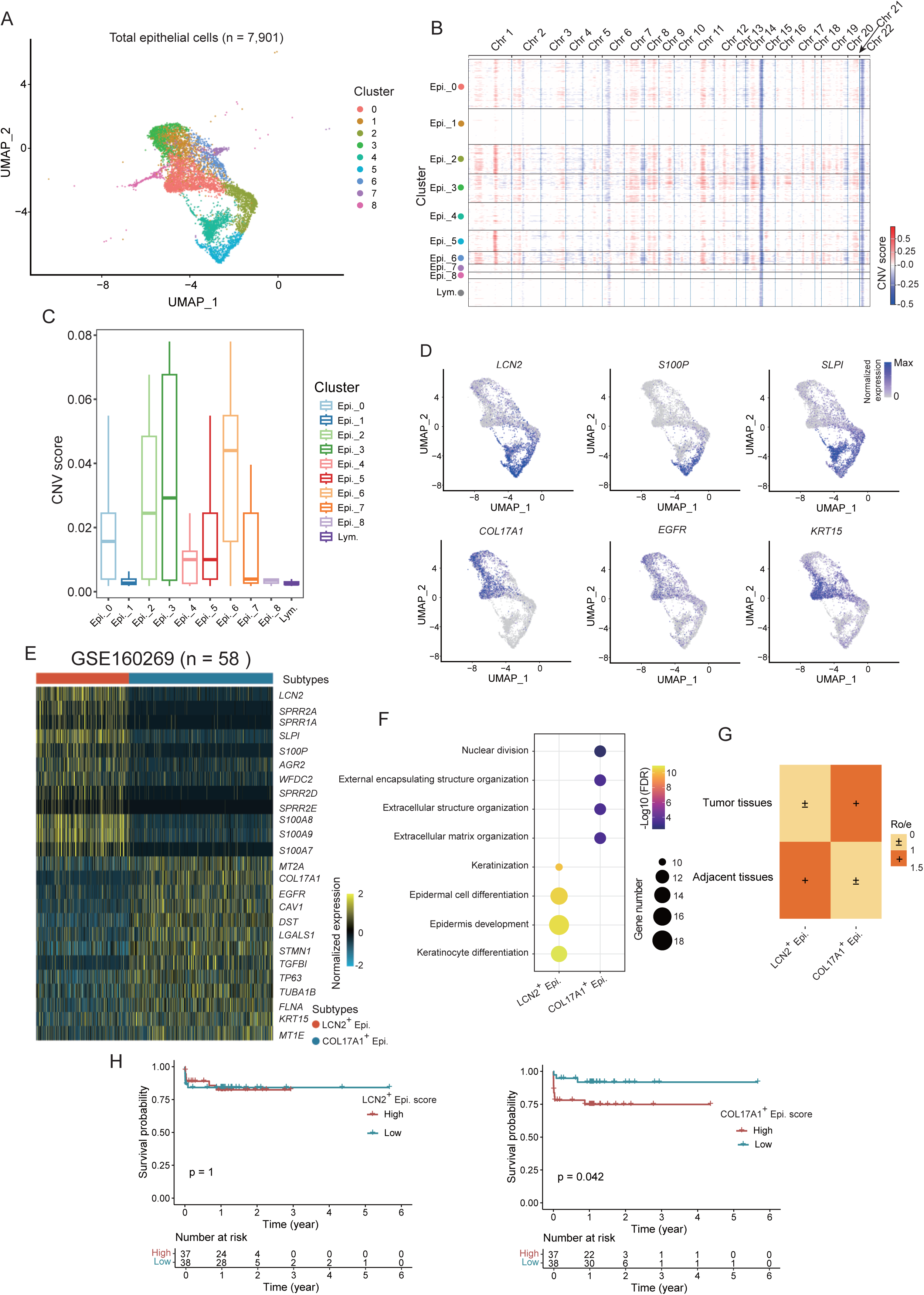
Epithelial cell clustering and functional analysis. **A.** UMAP plots of total epithelial cells labeled by seurat clusters. **B.** Heatmap showing the genomic CNV scores of nine epithelial cell clusters. **C.** boxplots showing the mean CNV scores across epithelial subpopulations. **D.** Feature plots of represented marker genes *LCN2*, *S100P*, *SLPI*, *KRT15*, *COL17A1*, and *EGFR*. **E.** Heatmap showing the expression of 26 signature genes in epithelial cells from GSE160269 dataset (n = 58). **F.** Dot plots showing enriched GO terms for the top 100 upregulated genes in each epithelial subpopulation. **G.** Heatmap showing cell preference distribution of *LCN2*^+^ or *COL17A1*^+^ epithelial cells in paired samples (n = 3), with Ro/e < 1 indicating nontumor-preferential distribution and Ro/e > 1 indicating tumor-preferential distribution. **H.** Kaplan–Meier survival curve for *LCN2*^+^ or *COL17A1*^+^ epithelial cell signature score in a TCGA-ESCC cohort (n = 75). Statistical significance is set at *P* < 0.05 (Wilcoxon test).

**Figure S6.**
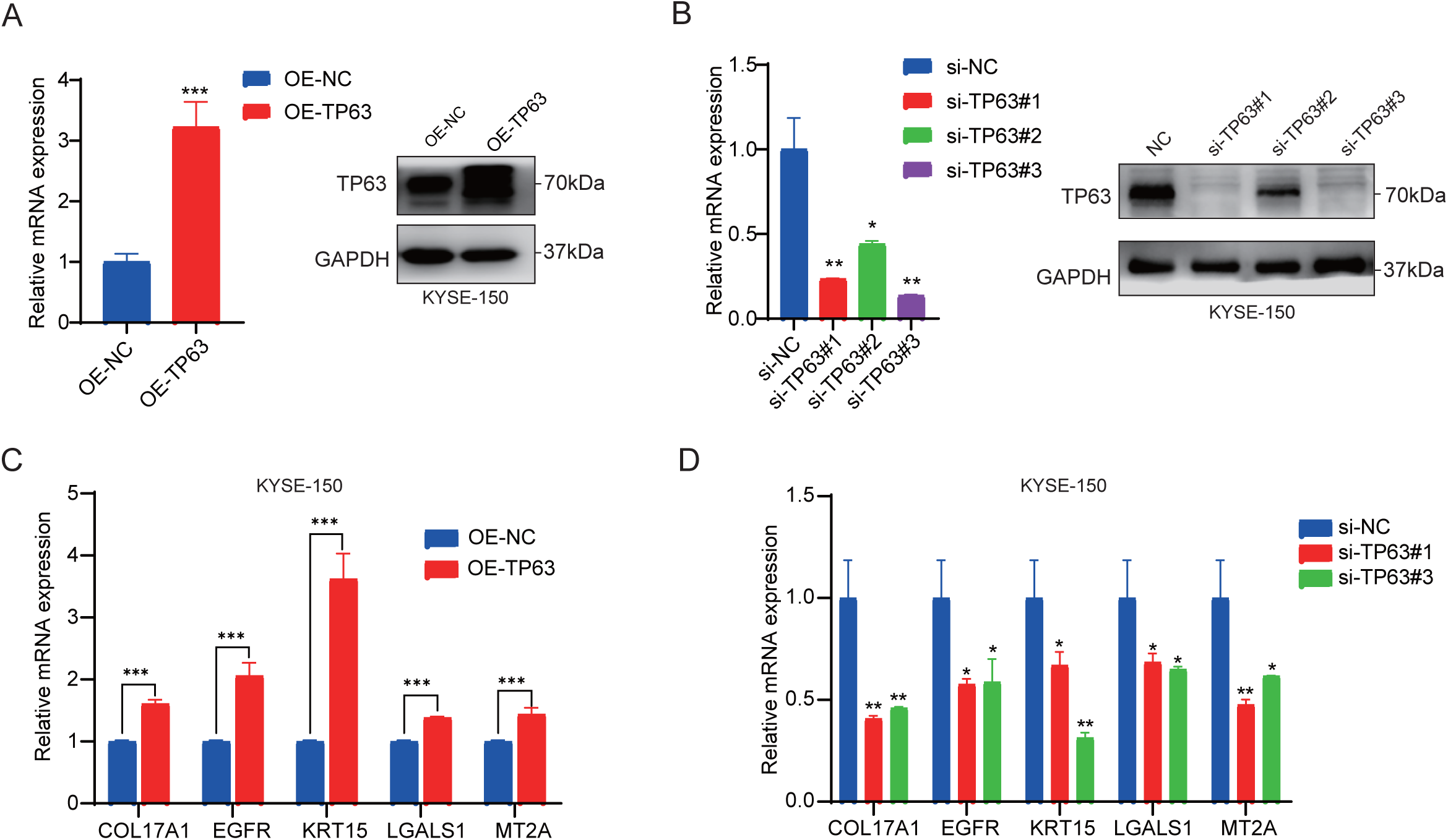
Validation of *TP63* overexpression and knockdown efficiency in KYSE-150. **A.** qPCR and WB analysis of *TP63* expression levels in KYSE-150 cells transfected with *TP63*-overexpressing lentivirus after 10 days. **B.** qPCR and WB analysis of *TP63* expression levels in KYSE-150 cells transfected with si-*TP63* after 48 h. qPCR analysis of genes *COL17A1*, *MT2A*, *EGFR*, *KRT15*, *LGALS1* after *TP63* overexpression (**C**) or knockdown (**D**) in KYSE-150 cells. ns, *P* is not significant; *, *P* < 0.05; **, *P* < 0.01; ***, *P* < 0.001. Statistical significance was calculated by Student’s *t*-test, mean ± SD. qPCR, quantitative real-time polymerase chain reaction.

**Figure S7.**
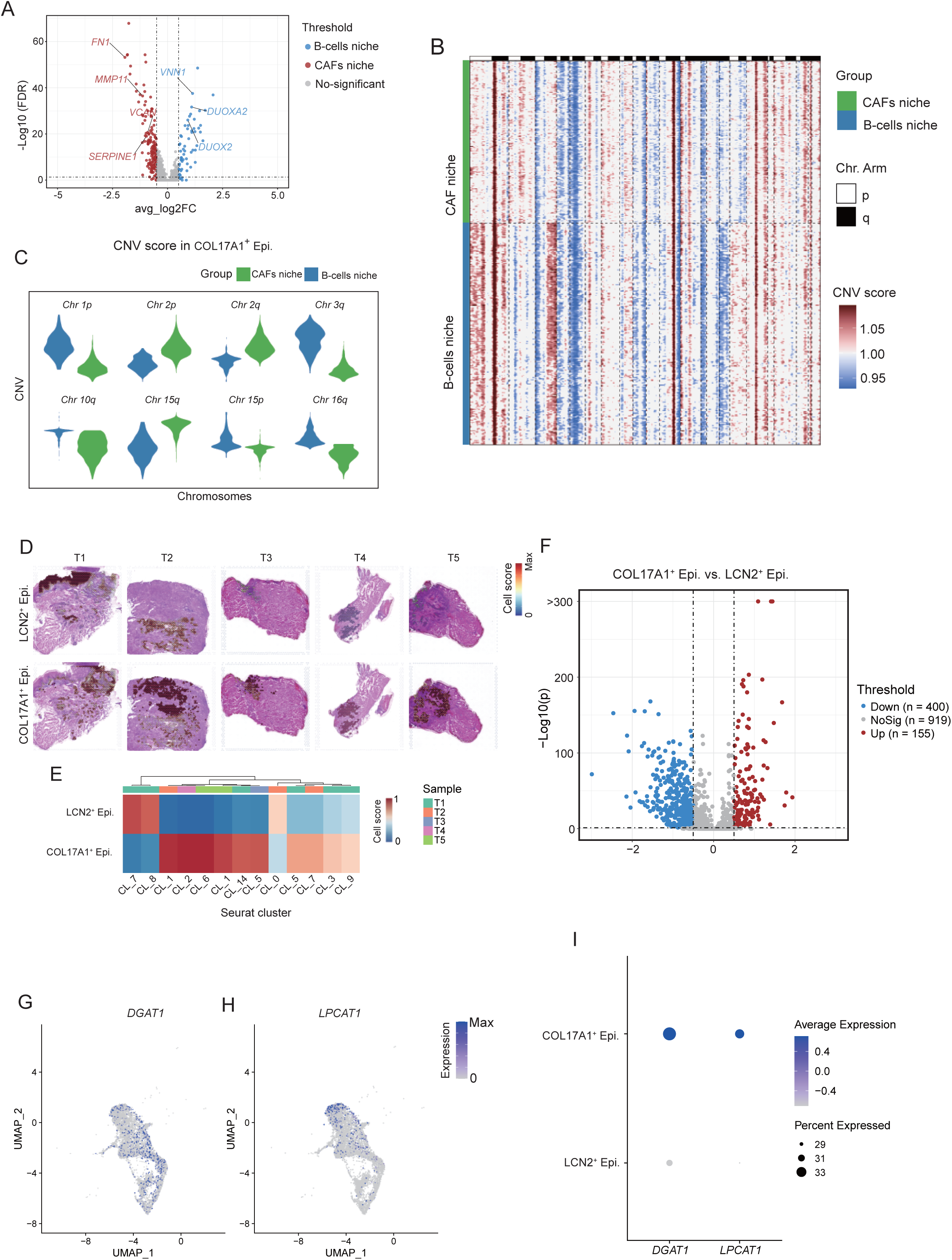
Spatial characteristics of epithelial subpopulations. **A.** Volcano plots showing differentially expressed genes for *COL17A1*^+^ epithelial cells between B cell and CAF niches in tumor sample 1. *P* < 0.05 and log_2_-fold change > 0.5 were considered as a statistically significant difference. **B.** Heatmap showing the genomic CNV scores of *COL17A1*^+^ epithelial cells in B cell or CAF niches. **C.** Violin plots showing the CNV score changes of representative chromosomes between B cell and CAF niches. **D.** Spatial feature plots showing the abundance of epithelial subpopulations in epithelial region across five tumor samples. **E.** Heatmap displaying the average abundance of epithelial subpopulations in seurat clusters across five tumor samples. **F.** Volcano plots showing differentially enriched metabolites between *LCN2*^+^ and *COL17A1*^+^ epithelial cells. *P* < 0.05 and log_2_-fold change > 0.5 are considered as a statistically significant difference. Feature plots of *DGAT1* (**G**) and *LPACAT1* (**H**) for epithelial subpopulations in scRNA-seq data. **I.** Dot plots showing the mRNA expression of *DGAT1* and *LPACAT1* in epithelial subpopulations from ST data.

**Figure S8.**
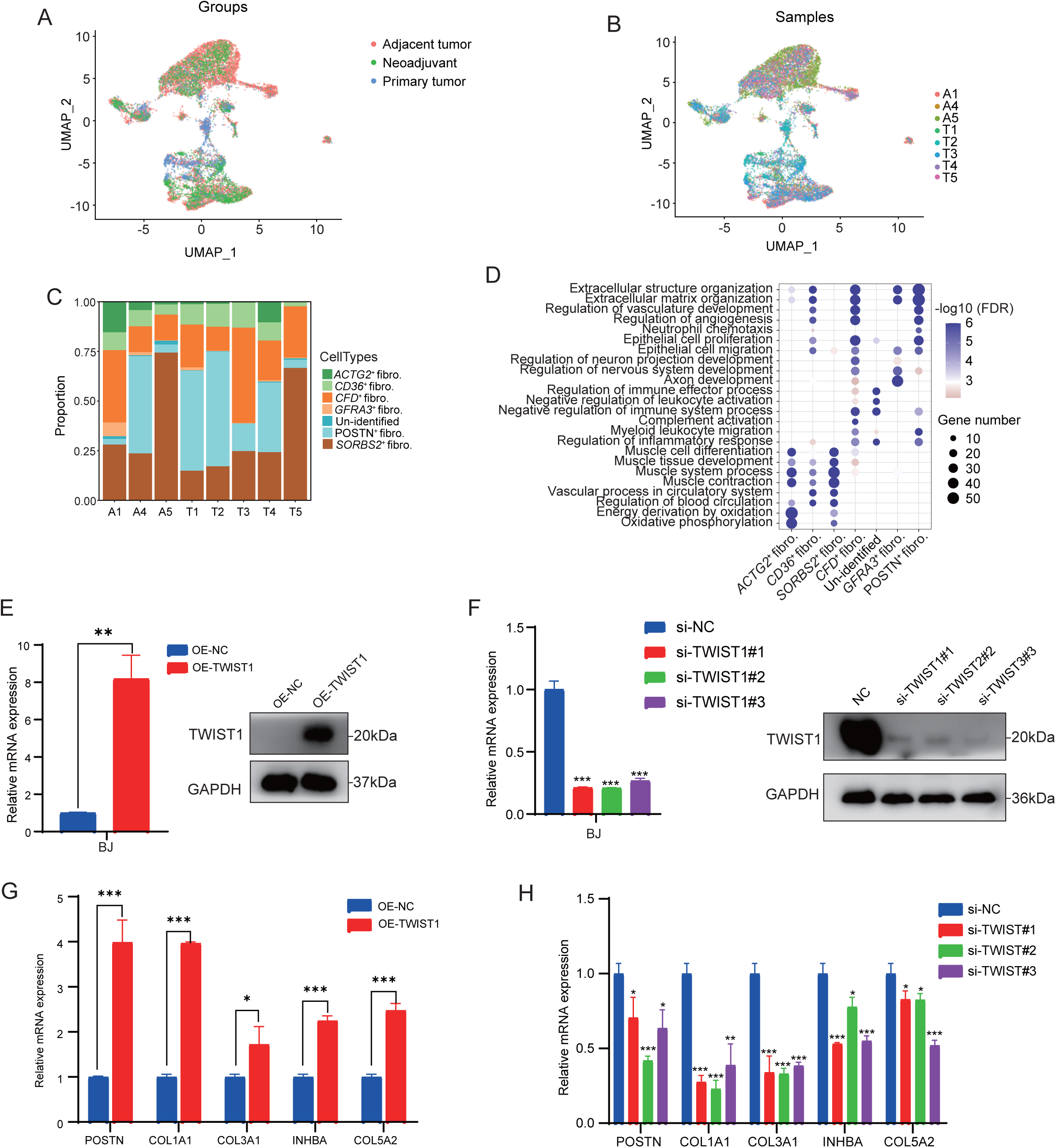
Validation of *TWIST1* overexpression and knockdown efficiency in BJ cells. UMAP plots of total fibroblasts labeled by groups (**A**) and samples (**B**). **C.** Bar plots showing fibroblast subcluster proportions across eight samples. **D.** GO terms of top 100 specifically expressed genes for each fibroblast subpopulation. **E.** qPCR and WB analysis of *TWIST1* expression levels in BJ cells transfected with *TWIST1* overexpression lentivirus after 10 days. **F.** qPCR and WB analysis of *TWIST1* expression levels in BJ cells transfected with si-*TWIST1* after 48 h. qPCR analysis of genes *POSTN*, *COL1A1*, *COL3A1*, *INHBA*, *COL5A2* after *TWIST1* overexpression (**G**) or knockdown (**H**) in BJ cells. *, *P* < 0.05; **, *P* < 0.01; ***, *P* < 0.001. Statistical significance was calculated by Student’s *t*-test, mean ± SD.

**Figure S9.**
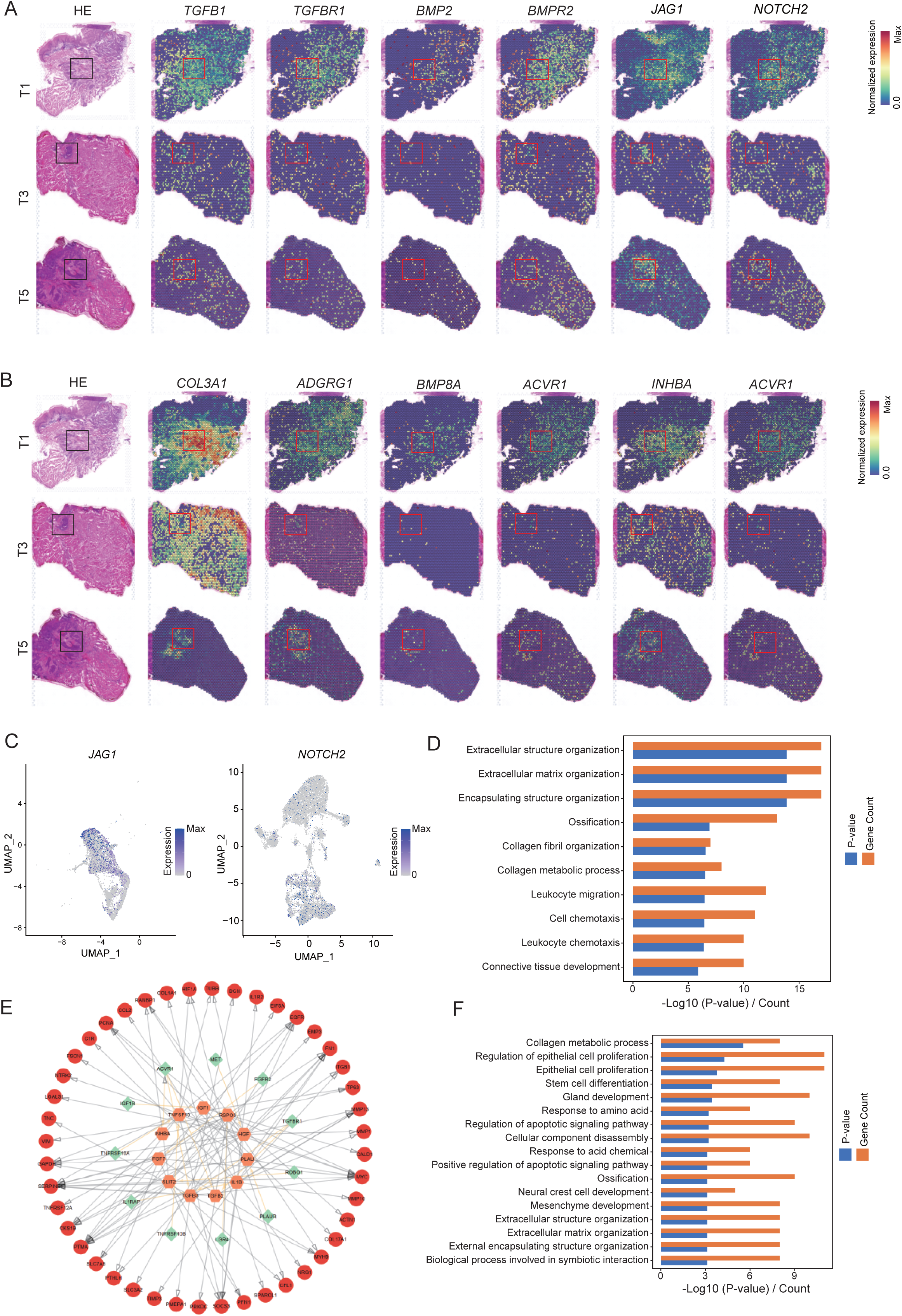
Representative cell–cell communication between tumor cells and fibroblasts. **A.** H&E staining overlays mRNA expression for selected L-R pairs (*COL17A1*^+^ epithelial cells to *POSTN*^+^/*CFD*^+^ fibroblasts) across tumor sample 1,3, and 5. **B.** H&E staining overlays mRNA expression for selected L-R pairs (*POSTN*^+^ fibroblasts to *COL17A1*^+^ epithelial cells) across tumor sample 1,3, and 5. **C.** Feature plots of scRNA-seq expression for *JAG1* and *NOTCH2*. **E.** The network of top active ligands from *POSTN*^+^ fibroblasts, matched receptors and matched target genes from *COL17A1*^+^ epithelial cells. Red represents ligands; orange represents receptors; green represents target genes. Represented GO terms of target genes expressed by *POSTN*^+^ fibroblasts (**D**) or *COL17A1*^+^ epithelial cells (**F**).

**Figure S10.**
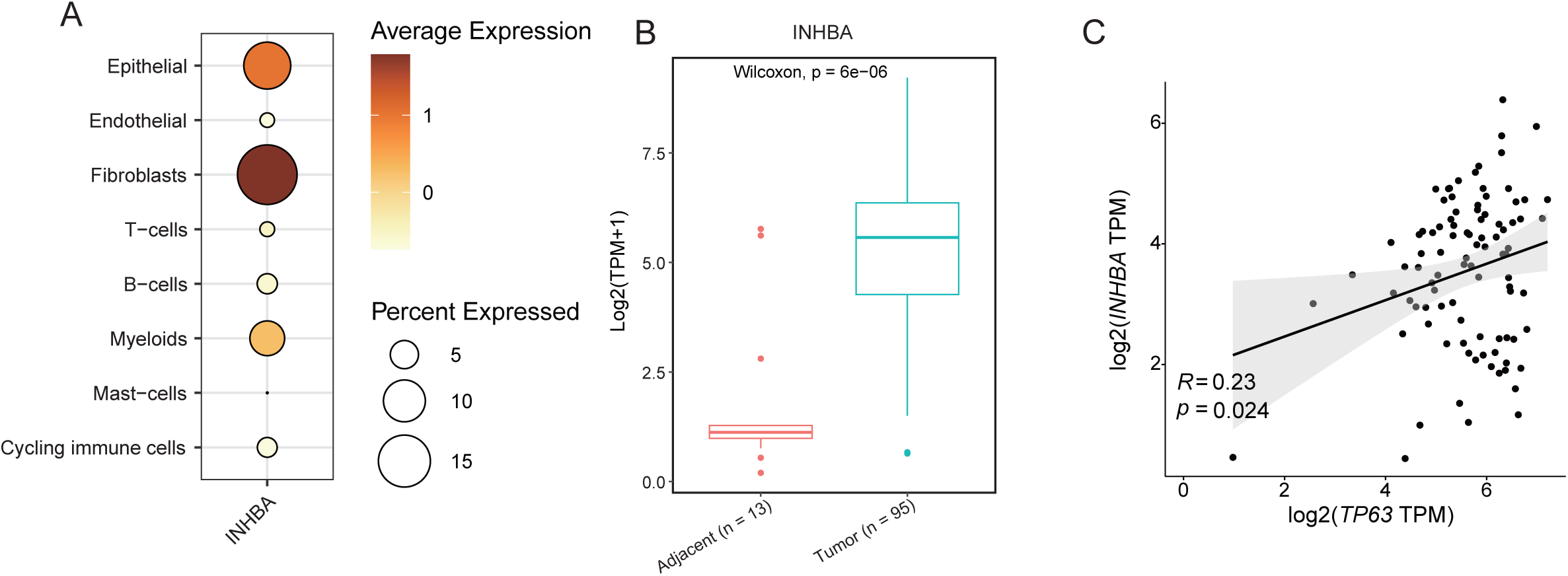
Analysis of *INHBA* expression levels and prognosis in public database. **A.** Dot plots showing the mRNA expression of *INHBA* across eight major cell types in scRNA-seq data. **B.** Box plots showing *INHBA* expression between ESCC (n = 95) and normal tissues (n = 13) in TCGA. **C.** Scatter plots showing the correlation between *INHBA* and *TP63* mRNA expression levels in TCGA-ESCC dataset (n = 95).

## Notes

### Competing Interest Statement

The authors have declared no competing interest.

