## Supplemental Table S1 for "A Single-cell and Spatially Resolved Cell Atlas of Human Esophageal Squamous Cell Carcinoma"

| Basic clinical information of patients included in this study | **Pathological grading** | II | II-III | II | II | Not reported |
| --- | --- | --- | --- | --- | --- | --- |
|  | **Neoadjuvent chemotherapy** | Karelizumab+cisplatin+paclitaxel | no | Karelizumab+cisplatin+paclitaxel | no | Karelizumab+cisplatin+paclitaxel |
|  | **Matched-adjacent cancerous tissues** | Yes | no | no | Yes | no |
|  | **Histological type** | Squamous cell carcinoma | Squamous cell carcinoma | Squamous cell carcinoma | Squamous cell carcinoma | Squamous cell carcinoma |
|  | **TNM** | pT3N1MX | Not reported | pT1aN0MX | pT3N1MX | T3-T4 |
|  | **Diagnosis** | ESCC | ESCC | ESCC | ESCC | ESCC |
|  | **Sex** | Male | Male | Female | Female | Male |
|  | **Age（years）** | 67 | 84 | 65 | 68 | 66 |
|  | **Patient ID** | 1 | 2 | 3 | 4 | 5 |
